# Functional proteomics identifies targetable cancer-associated fibroblast programs in head and neck cancer

**DOI:** 10.64898/2026.08.18.745234

**Authors:** Llara Prieto-Fernández, Ana Martínez-Carrillo, Lucas de Villalaín, Alejandra García-Torre, Beatriz de Luxán-Delgado, Francisco Hermida-Prado, Inmaculada Navarro-Lérida, Catalina Ribas, Ramón García-Escudero, Juan P. Rodrigo, Juan C. de Vicente, Tania Rodríguez-Santamarta, Juana M. García-Pedrero, Saúl Álvarez-Teijeiro

## Abstract

Head and neck squamous cell carcinoma (HNSCC) remains clinically challenging, with limited molecularly targeted options and a strong dependence on the tumor microenvironment. Cancer-associated fibroblasts (CAFs) are major stromal regulators that shape tumor progression, extracellular matrix remodeling, invasion, and therapeutic response. However, how CAF heterogeneity and plasticity translate into distinct tumor-promoting functions and targetable vulnerabilities remains insufficiently defined. Here, we integrated patient-matched primary CAFs and normal fibroblasts with 3D functional assays, tumor–stroma co-culture models, quantitative extracellular matrix analysis, whole-proteome profiling, and pharmacological perturbation. Primary fibroblast populations displayed marked interpatient heterogeneity and context-dependent plasticity in invasion, contractility, and responsiveness to tumor-derived signals, whereas enhanced fibronectin-rich matrix deposition and disorganization emerged as a conserved CAF-associated feature. Both normal fibroblasts and CAFs promoted HNSCC cell invasion in a population-dependent manner, whereas CAFs consistently induced less compact and more dispersed tumor nest architectures. Integrative functional analyses identified distinct CAF phenotypes characterized by either invasive and matrix-remodeling activity or high responsiveness to tumor-derived cues. Proteomic profiling revealed recurrent enrichment of adhesion, cytoskeletal, and extracellular matrix programs and guided the selection of pharmacological inhibitors aimed at modulating specific CAF-mediated pro-tumoral functions. Pharmacological targeting selectively altered these functions: CHI3L1 inhibition disrupted fibronectin matrix deposition, broad phosphodiesterase inhibition increased matrix alignment, and FZD7 inhibition consistently blocked tumor-induced CAF invasion across all tested populations. These findings define functionally distinct and pharmacologically targetable CAF programs in HNSCC and support stromal-directed interventions as a rational component of future combination treatment strategies.

## INTRODUCTION

Head and neck squamous cell carcinoma (HNSCC) is a heterogeneous malignancy of the upper aerodigestive tract and the seventh most common cancer worldwide (1,2). Despite advances in surgery, radiotherapy, chemotherapy and immunotherapy, outcomes remain poor in patients with advanced disease, for whom 5-year overall survival is approximately 40% (3,4). The limited availability of effective molecularly targeted therapies highlights the need to identify additional tumor-promoting mechanisms and therapeutic vulnerabilities.

Tumor progression is shaped not only by cancer cell-intrinsic alterations but also by dynamic interactions with the tumor microenvironment (TME). Cancer-associated fibroblasts (CAFs) are major stromal regulators of tumor growth, invasion, extracellular matrix (ECM) remodeling, therapeutic response and immune modulation in HNSCC (5,6). CAFs comprise heterogeneous and highly plastic populations with context-dependent functions ranging from tumor promotion to tumor restraint, and their phenotypic composition can evolve during cancer progression (7–10).

Although molecular and transcriptomic studies have extensively characterized fibroblast diversity in normal tissues and cancer (11,12), how CAF heterogeneity and plasticity translate into distinct tumor-promoting functions remains insufficiently understood. The scarcity of patient-matched primary fibroblast models preserving interpatient variability has also limited the systematic analysis of CAF functions, tumor– stroma crosstalk and pharmacologically targetable stromal vulnerabilities. This knowledge gap hinders the development of strategies that selectively modulate tumor-promoting CAF activities while preserving homeostatic or tumor-restraining functions.

Here, we integrated patient-matched primary CAFs and normal fibroblasts with 3D functional assays, tumor–stroma co-culture models, proteomic profiling and pharmacological perturbation. This strategy enabled us to define functionally distinct CAF phenotypes and identify molecular vulnerabilities that can be targeted to selectively modulate CAF-mediated pro-tumoral functions in HNSCC.

## METHODS

### Patient characteristics and sample collection

Fresh and formalin-fixed paraffin-embedded (FFPE) tissue samples were obtained from HNSCC patients diagnosed and surgically treated at the Oral and Maxillofacial Surgery Department of the Hospital Universitario Central de Asturias (HUCA). Patient-matched primary populations of CAFs and NFs were isolated from tumor tissue and adjacent normal mucosa, respectively, in collaboration with the Principado de Asturias Biobank. Six patient-matched CAF-NF pairs were selected and used for further *in vitro* functional and molecular characterization. The clinicopathological characteristics of the HNSCC patients and the primary tumors of origin included in the study are summarized in **Supplementary Table S1**.

### Primary fibroblast isolation

Tissue samples were minced and incubated with 100 U/mL collagenase IV (Gibco) in Hanks’ Balanced Salt Solution (HBSS) supplemented with caspofungin (0.5 µg/mL), for 90 min at 37°C. Cell suspensions were filtered through a 40 µm cell strainer, centrifuged and seeded in a 6-well plate. After 30 min of incubation, the supernatant containing non-adhered cells was transferred to another well, exploiting the rapid adhesion of fibroblasts to enrich the initial cultures. Fibroblasts were subsequently expanded and cryopreserved.

### Cell lines and culture conditions

The following HNSCC-derived cell lines were used: FaDu (pharynx; ATCC HTB-43) and Cal27 (tongue; CRL-2095) were purchased from the American Type Culture Collection; UT-SCC40 (tongue, T3N0M0) was kindly provided by Dr. Reidar Grenman (Turku University, Finland); and HCA-LSC1 (larynx, T3N2cM0) was established in our laboratory. Six patient-matched primary fibroblast pairs were also used, including CAFs (CAF1, CAF2, CAF4, CAF7, CAF8 and CAF12) and their corresponding NFs (NF1, NF2, NF4, NF7, NF8 and NF12). All models originated from HPV-negative primary tumors. Cell cultures were routinely tested for mycoplasma using a PCR-based kit (Biotools). Cell line authentication was performed by DNA (STR) profiling at the SCT Core Facilities (University of Oviedo, Spain).

Cells were maintained in Dulbecco’s Modified Eagle Medium (DMEM) culture medium containing 4.5 g/L glucose and 2 mM L-glutamine (Corning), supplemented with 10% fetal bovine serum (FBS; Corning), 100 U/mL penicillin/streptomycin (Biowest), 20 mM HEPES (Biowest) and 100 mM non-essential amino acids (Biowest), hereafter referred to as complete DMEM. For specific experimental procedures, complete DMEM was used without FBS when indicated. Primary fibroblasts were additionally supplemented with 1% ITS (insulin-transferrin-selenium; Fisher). Cells were cultured in a humidified atmosphere with 5% CO_2_ at 37°C. Primary fibroblasts were used up to passage 10.

### Conditioned media production

HNSCC cells were seeded in T175 flasks and cultured to approximately 60% confluence. Cells were washed twice with phosphate-buffered saline (PBS) and maintained in serum-free DMEM for 72 h. Conditioned media (CM) were collected and centrifuged at 2,000 rpm for 10 min to remove cell debris, aliquoted and stored at -80°C until use.

### 3D spheroid invasion assays

3D collagen invasion assays were performed as previously described (13). Briefly, cells were suspended in complete DMEM containing 20% methylcellulose (Sigma) at a density of 80,000 cell/mL. Twenty-five microliter drops of cell suspension were pipetted onto a non-adhesive Petri dish and incubated in inverted position overnight at 37°C. The following day, each spheroid was transferred to a well of a 96-well plate and embedded in 110 μL of a 2.3 mg/mL type I collagen matrix (PureCol®, Advanced BioMatrix). After 90 min polymerization at 37°C, 100 μL of medium were added to each well. Fibroblast spheroid invasion assays were monitored for 20 h, whereas HNSCC cell spheroid and co-spheroid invasion assays were monitored for 24 h. Images at initial and final time points were acquired using a Leica DMi1 microscope (Leica Microsystems) for fibroblast spheroids. For co-spheroid experiments, cells were monitored using a Zeiss Cell Observer Live Imaging microscope (Zeiss) coupled with a CO_2_ and temperature-maintenance system for live cell tracking. Time-lapse images were acquired every 60 min for 24 h using a Zeiss AxioCam MRc camera (Zeiss). Spheroid area was measured using ImageJ software. The invasive area was calculated by normalizing the spheroid area at the final time point to the area at time 0, and expressed relative to the control condition when indicated.

For co-spheroids, fibroblasts and cancer cells were mixed at a 3:1 ratio. HNSCC cells were labelled with CellTracker^TM^ Green CMFDA (Fisher) at 3 μM for 30 min. For invasion analysis at 24 h, only the green fluorescent area corresponding to cancer cells was measured. In experiments using fibroblast-containing matrices, HNSCC spheroids were embedded in collagen matrices containing 20,000 fibroblasts per well.

### Collagen gel contraction – ECM remodeling assays

48-well plates were coated with 250 μL of 1% bovine serum albumin (BSA) in PBS and left overnight. Fibroblasts were then embedded at 80.000 cells per 100 μL of matrix in matrices composed of 4 mg/mL rat tail collagen I (Corning) and 2 mg/mL Gel ECM from Engelbreth-Holm-Swarm murine sarcoma (Sigma). A total of 120 μL of cell-matrix suspension was carefully seeded per well, and incubated for 1 h at 37°C, before adding 300 μL of culture medium. Images were acquired at 24 h using a GS-800 Calibrated Densitometer scanner (Bio-Rad). Total well and gel areas were measured using ImageJ software, and contraction was calculated as 100 × [(total area − gel area)/total area].

### ECM deposition/synthesis assays

12-mm-diameter glass coverslips were coated overnight with 1 μg/mL fibronectin in 24-well plates. The following day, 25,000 fibroblasts were seeded per well and cultured for 5 days to allow for ECM deposition and remodeling. Cells were then fixed and processed for Immunofluorescence.

Quantitative analysis of ECM fluorescence images was performed using ImageJ software. Fibronectin fiber anisotropy was assessed using the *FibrilTool* (14) ImageJ plug-in, which generates anisotropy scores ranging from 0 (indicating disorganized matrixes with randomly oriented fibers) to 1 (representing organized matrices with uniformly aligned fibers). Fiber orientation was further analyzed using the *Directionality* tool within ImageJ (15). This analysis generates histograms representing the distribution of fibers across different orientations. Isotropic matrices lacking a preferential direction are expected to produce flat histograms, whereas histograms from ordered matrices display a peak corresponding to the predominant fiber orientation. The highest peak in the orientation histogram is fitted by a Gaussian function, and the following parameters were extracted: *Dispersion (°)*, indicating the deviation from the Gaussian fit and reflecting the spread of fiber orientations (lower dispersion values are expected in ordered matrices); *Amount*, representing the percentage of fibers aligned in the preferred orientation; and *Goodness*, measuring the quality of the Gaussian fit.

To obtain a composite metric integrating these three parameters, in which higher values correspond to more ordered matrices, the *Directionality Index* was defined as:

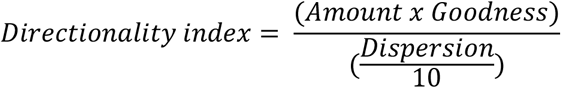

### Co-culture nest formation assays

12-mm-diameter glass coverslips were coated overnight with 1 μg/mL fibronectin in 24-well plates. The following day, 25,000 fibroblasts and 25,000 HNSCC cells were seeded per well and co-cultured for 3 days to allow nest formation. Cultures were then fixed and processed for Immunofluorescence. Tumor nest circularity was quantified using the ImageJ *Circularity* tool (16).

### Immunofluorescence

Cells were fixed with 4% paraformaldehyde in PBS for 15 min at room temperature (RT), permeabilized with 0.5% Triton X-100 in PBS for 10 min at RT, and blocked with 3% BSA in PBS containing 1% Tween-20 for 30 min at RT. Primary antibodies were incubated overnight (**Supplementary Table S2**), followed by fluorescence secondary antibodies for 1 h (**Supplementary Table S2**) and, where indicated, Phalloidin 546 (Fisher; 1:40 dilution) and DAPI (Fisher; 1:1000 dilution). Coverslips were mounted onto glass slides using Dako Fluorescence Mounting Medium (Agilent) and fluorescence images were acquired using a Zeiss Cell Observer microscope equipped with a Zeiss AxioCam MRc camera (Zeiss).

### Proliferation assays

Fibroblasts were seeded at 3,000 cells per well in 96-well plates. Proliferation was measured at baseline and after 96 h using the CyQuant Cell Proliferation Assay Kit (Invitrogen) following the manufacturer’s instructions. Fluorescence was measured at 480/520 nm using a Synergy HT microplate reader (BioTek). Values at 96 h were normalized to baseline and expressed relative to the control condition.

### Pharmacological inhibitors

The pharmacological inhibitors used in this study are listed in **Supplementary Table S3**. Drug concentrations and treatment conditions are specified within each assay.

### Drug cell viability assays

Fibroblasts were seeded at 2,500 cells per well in 96-well plates and treated with the indicated compounds for 24 h, 48 h, 72 h or 5 days. At each endpoint, cell viability was assessed using the CellTiter 96 Aqueous One Solution MTS Assay (Promega). Absorbance was measured at 490 nm using a Synergy HT plate reader (BioTek). values were normalized to the baseline and expressed relative to control condition.

### LC-MS/MS proteomics

Fibroblasts were seeded in collagen I-coated (40 μg/mL, PureCol®, Advanced BioMatrix) 100 mm culture dishes and grown for 48 h. Cells were scraped on ice and protein lysates were collected using a lysis buffer containing 50 mM Tris-HCl (pH 7.5), 150 mM NaCl, 10% glycerol, 1% NP-40 and Halt^TM^ Protease & Phosphatase Single-Use Inhibitor cocktail (Thermo Fisher Scientific). Four independent biological replicates were collected from each fibroblast population. Proteins were precipitated with cold acetone overnight, resuspended in 0.2% RapiGest in 50 mM ammonium bicarbonate, and sonicated. Protein concentration was determined using a Qubit4 fluorometer (Invitrogen). A total of 40 μg of protein were used for further digestion. For disulfide bond reduction and alkylation, samples were sequentially incubated with 50 mM DTT in 50 mM ammonium bicarbonate for 30 min at 60°C, followed by 100 mM iodoacetamide for 30 min at RT in the dark. For digestion, trypsin was added at a 1:40 enzyme:protein ratio and incubated for 2h at 37°C, followed by an additional overnight incubation step. The reaction was stopped by adding 10% trifluoroacetic acid (TFA) and incubating for 1h at 37°C. Acetonitrile (ACN) was then added and samples were centrifuged. Supernatants were transferred to new tubes and stored at -20°C until mass spectrometer injection.

LC-MS/MS procedures were performed at the Proteomics Unit of ISPA-FINBA (Oviedo, Spain). For each sample, 400 ng of digested proteins were loaded onto Evotips (Evosep) and analyzed in a hybrid Q-TOF mass spectrometer (ZenoTOF 7600, Sciex) coupled to an Evosep One liquid chromatography system (Evosep). One randomly selected sample was run in triplicate to estimate the coefficients of variation (CVs) for each protein. Peptide digests were separated using the 30 samples per day Evosep program (44 min total run time) with water (solvent A) and acetonitrile (solvent B), both containing 0.1% formic acid. Column temperature was set at 40 °C. Peptide ionization was performed using an Optiflow electrospray ion source (Sciex) equipped with a low-micro electrode, applying a voltage of 4500 V and 100 °C. ZenoSWATH data-independent acquisition (DIA) (17) was used as acquisition method. In this approach, data are acquired in repeated acquisition cycles consisting of one TOF MS scan (350-1250 m/z, 50 ms accumulation time), followed by 85 MS/MS scans using variable Q1 isolation windows covering the 349.5-1247 m/z range. MS/MS spectra were acquired over the 230-1400 m/z range with a 20 ms accumulation time, with Zeno pulsing activated and dynamic collision energy applied. Automatic calibration of TOF MS and MS/MS was performed after each sample using the X500 ESI Positive Calibration Solution (Sciex) through the calibrant delivery system of the mass spectrometer.

The ZenoSWATH runs were processed with DIA-NN v1.8.1 software (18) following the library-free workflow according to the instructions from the authors. First, DIA-NN built an *in silico*-predicted spectral library from the SwissProt human protein database, including isoforms (42,332 entries). The ZenoSWATH runs were then analysed against this predicted library. The main parameters used in DIA-NN were: 0 missed cleavages; N-terminal Met excision and Cys carbamidomethylation as fixed modifications; 2 to 5 precursor charge range; 350 to 1,500 precursor m/z range; 200 to 1,800 fragment ion m/z range; match-between-runs enabled; neural network classifier in double-pass mode; quantification strategy set to robust LC (high precision); and RT-dependent cross-run normalization. Protein groups were identified and quantified using only proteotypic peptides, applying a 1% false discovery rate (FDR) for both protein groups and precursors.

### Proteomics data analysis

Protein intensity matrix generated by DIA-NN software was further analyzed using the R environment (v4.4.1) (19). Proteins with CV <20% were selected for further analysis. Intensity values were log2-transformed and proteins with ≥60% missing values (NA) were excluded. Data were normalized by applying the quantile normalization method using the preprocessCore R package (v1.66.0) (20). Missing value imputation was performed using the quantile regression imputation of left-censored data (QRILC) approach implemented in the imputeLCMD R package (v2.1) (21). Differential protein expression analysis was performed using the limma R package (v3.60.4) (22) to fit our dataset to a linear model and compute empirical Bayes statistics. In this model, differences in protein expression were estimated based on pairwise comparisons of group means. Proteins with adjusted *p*-values < 0.05 were considered statistically significant. Subsequent enrichment analyses were performed using the clusterProfiler (v4.12.2) (23), org.Hs.eg.db (v3.19.1) (24), ReactomePA (v1.48.0) (25) and reactome.db (v1.88) (26) R packages. For Gene Ontology (GO) over-representation analysis (ORA) and Reactome ORA, proteins with adjusted *p*-values < 0.05 were used as input, and the expression matrix of the 5,762 identified proteins was used as *universe* for GO ORA analysis.

Weighted gene co-expression network analysis (WGCNA) was performed on the normalized and imputed protein intensity matrix, considering proteins that were differentially expressed in at least one CAF-NF comparison. The analysis was conducted using the WGCNA R package (v1.72.5) (27), following the pipeline developed by Wu *et al*. (28). A correlation cut-off of 0.85 and a soft-thresholding power of 6 were selected to construct the weighted correlation and adjacency matrices according to the approximate scale-free topology criterion. From the adjacency matrix, the topology overlap matrix (TOM) and corresponding distances were calculated and used for hierarchical clustering (average linkage method) based on TOM distance. Optimal clusters (modules) were determined by dynamic tree cutting algorithm (minimum module size = 10 proteins). Closely correlated modules were then merged using a cut height of 0.3.

Eigenproteins from each module were calculated as the first principal component (PC) of the expression matrix for each respective module via singular value decomposition. Eigenprotein-based module memberships (kME) were then computed to assess the correlation between each protein expression profile and the corresponding module eigenprotein. Proteins with higher kME values were considered hub proteins, displaying strong correlation with the eigenprotein from their module. Finally, Pearson correlations between module eigenproteins and biological groups (i.e., each fibroblast subpopulation) were calculated to identify trait-associated modules.

### Western Blot

Fibroblasts were seeded in collagen I-coated (40 μg/mL, PureCol®, Advanced BioMatrix) 100 mm culture dishes and grown for 96 h. In the experiments involving CM treatment, fibroblasts were treated with CM for 72 h. Protein lysates were extracted using Laemmli buffer supplemented with 1 mM dithiothreitol (DTT) (Sigma) and Halt^TM^ Protease & Phosphatase Single-Use Inhibitor cocktail (Thermo Fisher Scientific), and then briefly sonicated. Protein concentration was determined using the Pierce BCA Protein Assay kit (Thermo Fisher Scientific). Equal amounts of protein (40 μg of each protein extract supplemented with β-mercaptoethanol and boiled) were loaded onto SurePAGE™ Bis-Tris 4-12% precast polyacrylamide gels (GenScript) and subsequently transferred to nitrocellulose membranes using the Trans-Blot® Turbo^TM^ Transfer System (BioRad). Membranes were blocked with 5% BSA for 1 h and incubated overnight with the correspondent primary antibodies (**Supplementary Table S4**). Fluorescence-conjugated secondary antibodies were used at 1:10,000 dilution for detection (**Supplementary Table S4**). Membranes were scanned using the Odyssey Fc Dual-Mode Imaging System (LICORbio) in the 700nm (red) and 800nm (green) channels. Densitometric analysis was performed using Image Studio Lite software (LICORbio). Protein expression levels were normalized to GAPDH as loading control.

### Statistical analysis

Statistical analyses of *in vitro* assays were performed using GraphPad Prism (v8.0.2). Unpaired *t*-test, one- and two-way ANOVA and Dunnett’s or Tukey’s multiple comparisons tests were applied as indicated. Proteomic analyses were performed in R.

## RESULTS

### Primary stromal fibroblasts isolated from HNSCC patients retain high functional heterogeneity

Twelve primary fibroblast populations comprising six patient-matched CAFs-NFs pairs (1, 2, 4, 7, 8, and 12), were isolated from HNSCC tumors and adjacent normal tissues. CAF1, CAF2, and CAF4 showed increased growth compared with their matched NFs at 96 h, whereas CAF7 and CAF8 displayed reduced proliferation, and CAF12 showed no significant difference (**Figure 1A**).

**Figure 1.**
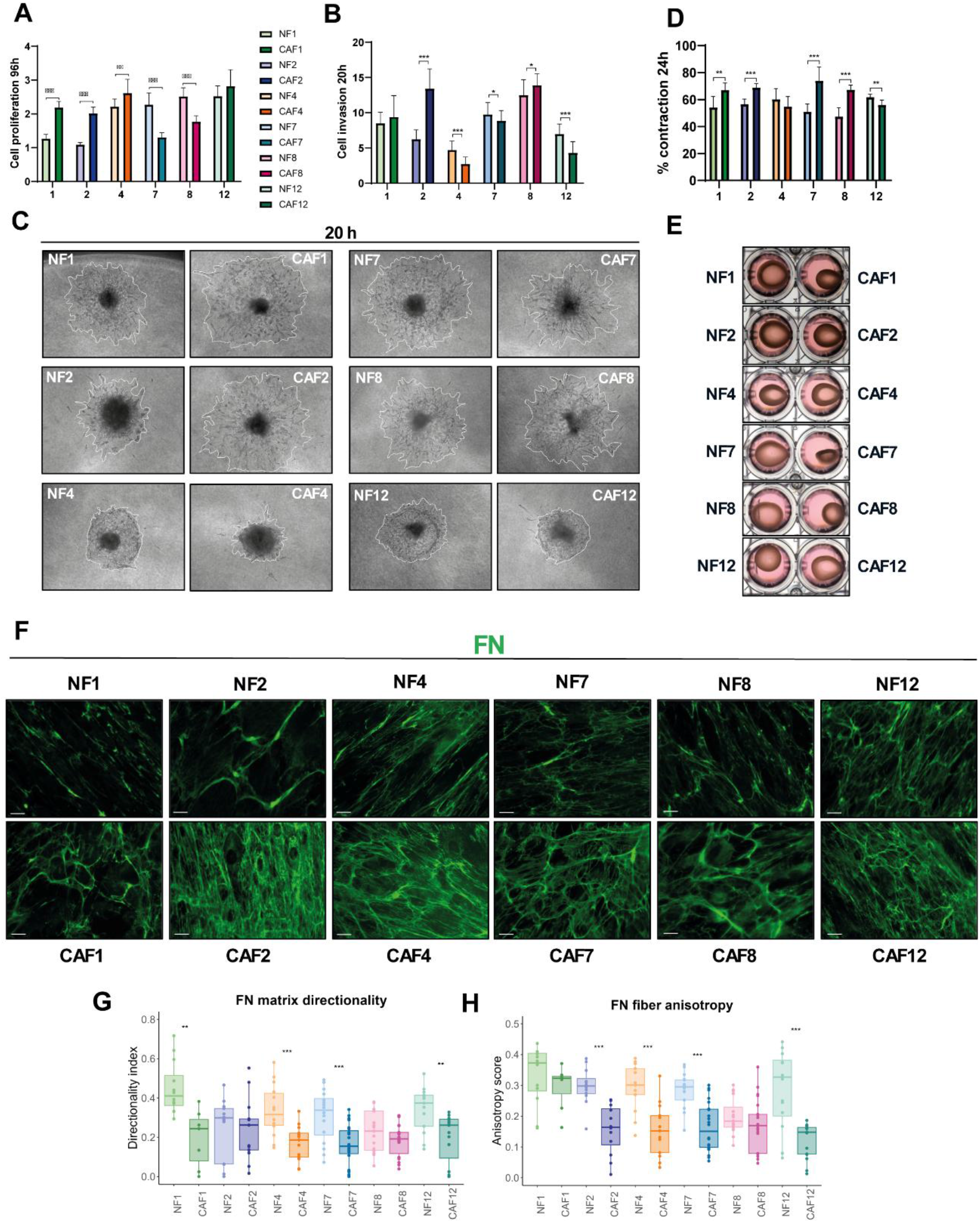
Functional diversity of patient-matched primary CAF and NF fibroblast populations. **A.** Fibroblast proliferation measured by CyQuant after 96 h, data normalized to day 0. n = 3 independent experiments, performed in triplicate. **B.** Fibroblast 3D invasion at 20 h, normalized to initial spheroid area at 0 h. n ≥ 3 independent experiments, in triplicate. **C.** Representative end-point images of fibroblast spheroid invasion at 20 h for each fibroblast population. The final invading area is highlighted in white. **D.** Percentage of collagen contraction after 24 h. n = 3 independent experiments, in duplicate. **E.** Representative end-point images of 3D collagen contraction assays at 24 h for each fibroblast population. Data are represented as mean ± SD. Statistical significance for each CAF *vs*. NF pair was calculated using unpaired *t*-test. \**p* < 0.05, ** *p* < 0.01 and *** *p* < 0.001. **F**. Representative immunofluorescence images of fibroblast-derived fibronectin ECM after 5 days in culture. n = 3 independent experiments. Green = fibronectin (FN). Scale bar = 20 µm. **G.** Directionality index of fibronectin matrices. **H.** Anisotropy scores of fibronectin fibers. Each dot represents a single image measurement. Measurements were performed in at least four images per assay across 3 independent assays. Statistical significance for each CAF *vs*. NF pair was determined using unpaired *t*-test. \**p* < 0.05, ** *p* < 0.01 and *** *p* < 0.001.

Fibroblast invasive into 3D collagen matrices exhibited marked heterogeneity, with pairs 1, 2, 7 and 8 being highly invasive, whereas pairs 4 and 12 showed reduced invasiveness (**Figure 1B, C**). Within matched pairs, CAF2 and CAF8 were more invasive than their corresponding NFs, whereas CAF4, CAF7, and CAF12 showed lower invasion. No significant differences were observed between CAF1 and NF1, although CAF1 showed a trend towards increased invasion (**Figure 1B, C**). Matrix contraction was likewise heterogeneous across populations, with CAF1, CAF2, CAF7, and CAF8 displaying greater contractile capacity than their matched NFs, whereas CAF12 contracted less than NF12 and CAF4 showed a similar, although non-significant, downward trend. Overall, this pattern broadly mirrored fibroblast invasion, with the most invasive populations also tending to display greater matrix-contractile activity (**Figure 1D, E**).

Analysis of ECM remodeling and deposition revealed a more consistent CAF-associated pattern. CAFs generated denser, patchier, and more disorganized fibronectin matrices than matched NFs, although the extent of this remodeling varied across populations and was most evident in CAF7 and CAF8 **(Figure 1F; Supplementary Figure S1).** Quantitative analysis confirmed reduced matrix directionality and anisotropy in most CAF-derived matrices, reaching statistical significance in pairs 1, 4, 7, and 12 **(Figure 1G, H; Supplementary Figure S2)**.

Overall, these results indicate that the primary fibroblast populations retain high functional heterogeneity under *in vitro* culture conditions. Interestingly, ECM remodeling and deposition were the only common features shared across all CAF populations, which consistently produced denser and more disorganized matrices.

### Crosstalk between HNSCC cells and primary stromal fibroblasts

To investigate the bidirectional tumor-stroma crosstalk, we combined conditioned-media experiments modeling paracrine signaling with 3D co-culture systems assessing direct tumor–fibroblast interactions. HNSCC-derived CM induced heterogeneous effects on fibroblast invasion, with UT-SCC40 and Cal27-derived CM consistently eliciting the strongest pro-invasive responses. Fibroblast sensitivity to these tumor-derived signals was also population dependent, with NF1, CAF1, CAF4, NF12, and CAF12 showing the largest and statistically significant increases in invasion **(Figure 2A)**. In 3D co-spheroid models, most fibroblast populations enhanced HCA-LSC1 and Cal27 invasion compared with tumor-only spheroids, pulling tumor cells and facilitating their spreading through the collagen matrix, although the magnitude of this effect varied across populations. In matched comparisons, CAF2 and CAF4 significantly increased HCA-LSC1 invasion relative to their corresponding NFs, whereas CAF4, NF8, and CAF12 showed significantly greater effects than their matched populations in Cal27 co-spheroids **(Figure 2B–D)**. Similarly, embedding Cal27 spheroids in fibroblast-containing collagen matrices increased tumor invasion in all conditions except NF1 **(Figure 2E).** Together, these findings demonstrate that both primary NFs and CAFs can potentiate HNSCC cell invasion, highlighting their functional plasticity as invasion-promoting stromal populations in response to tumor-derived cues.

**Figure 2.**
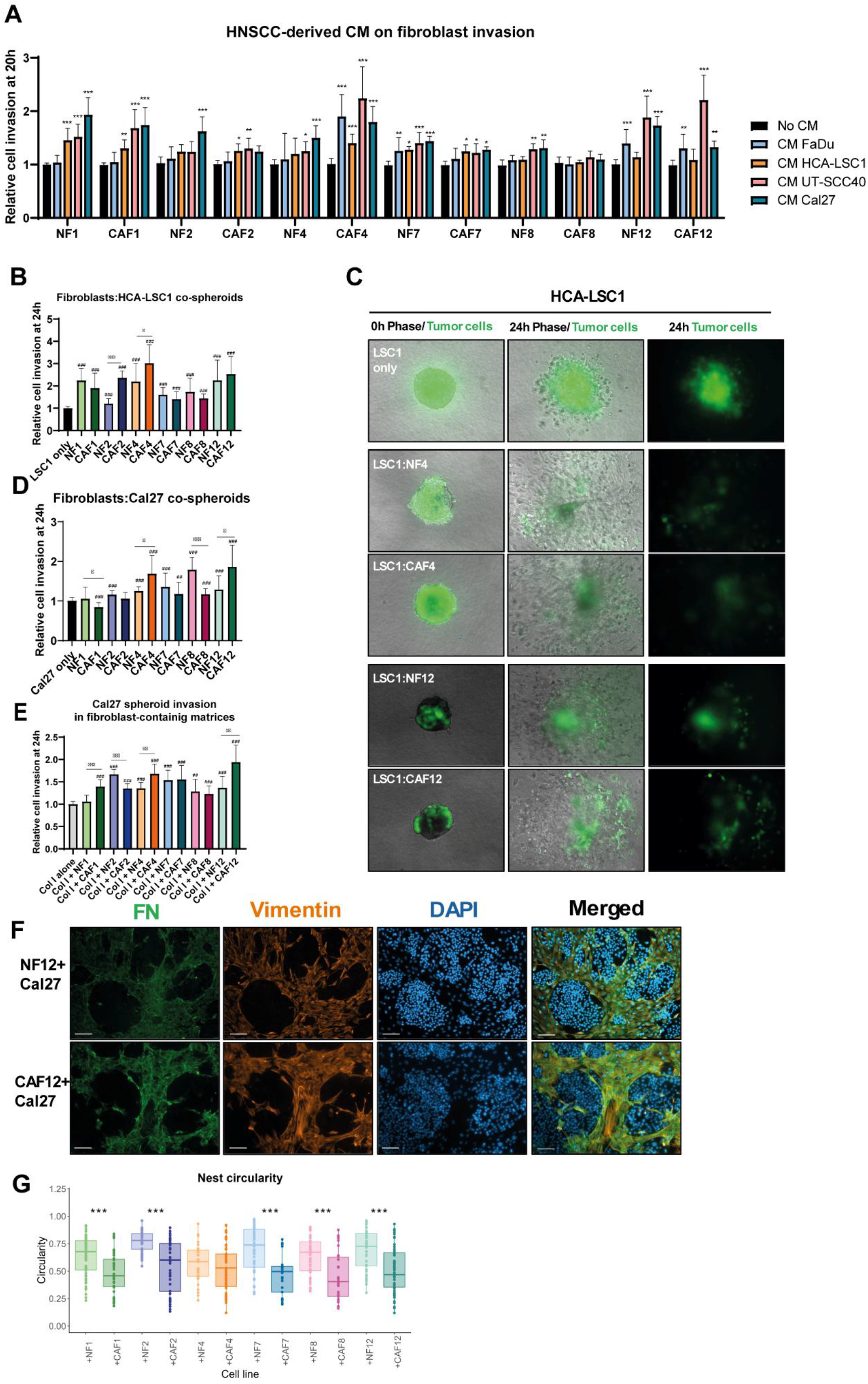
Bidirectional tumor-stroma crosstalk. **A.** Relative fibroblast 3D invasion at 20 h in response to HNSCC-derived conditioned media (CM), normalized to area at initial time (0 h) and relative to each control condition (No CM). n ≥ 3 independent experiments, performed in triplicate. Data are represented as mean ± SD. Statistical significance was calculated using Dunnett’s multiple comparisons test. \**p* < 0.05, ** *p* < 0.01 and *** *p* < 0.001. **B.** Co-spheroids of fibroblasts and HCA-LSC1 cells (3:1 ratio) in a 3D collagen matrix. Green-labelled tumor invading area was quantified after 24 h, normalized to area at initial time (0 h) and relative to control condition (LSC1 only). **C**. Representative images of HCA-LSC1 co-spheroid 3D invasion with fibroblast pairs 4 and 12. **D.** Co-spheroids of fibroblasts and Cal27 cells (3:1 ratio) in a 3D collagen matrix. Green-labelled tumor invading area was quantified after 24 h, normalized to area at initial time (0 h) and relative to control condition (Cal27 only). **E.** Cal27 spheroid invasion into a fibroblast-containing matrix at 24 h, normalized to area at initial time (0 h) and relative to control condition (Col I matrix without fibroblasts, i.e. “Col I alone”). Data is represented as mean ± SD. Statistical significance is indicated by # for comparisons with control condition, and by * for paired CAF-NF comparisons. * or # *p* < 0.05, ** or ## *p* < 0.01 and *** or ### *p* < 0.001. n ≥ 3 independent experiments in triplicate. **F.** Representative immunofluorescence images of co-cultured Cal27 cells with the indicated fibroblast population (proportion 1:1) after 72 h. Green = fibronectin (FN); blue = DAPI; orange= Vimentin. Scale bar = 100 µm. **G.** *Circularity* measures of tumor nests in the FN channel. Each dot represents an individual nest shape measurement. Only complete nests within an image were considered for analysis. n = 3 independent experiments, with at least three images per condition. Statistical significance was calculated for each CAF-NF comparison using unpaired *t*-test. \**p* < 0.05, ** *p* < 0.01 and *** *p* < 0.001.

As an additional approach to assess bidirectional tumor–stroma crosstalk, HNSCC tumor nest formation was analyzed in co-cultures with primary fibroblast populations. Compared with matched NFs, CAFs generated less compact and more interspersed Cal27 nests, with fibroblasts infiltrating tumor clusters. Circularity analysis confirmed more irregular tumor nest morphology in the presence of all CAF populations except CAF4 **(Figure 2F-G and Supplementary Figure S3)**.

Collectively, these findings demonstrate dynamic reciprocal communication between HNSCC cells and primary stromal fibroblasts across multiple 3D co-culture models, with marked interpatient heterogeneity. Despite this variability, CAFs generally showed stronger invasion-promoting effects than their matched NFs and consistently induced less compact and more dispersed tumor nest architectures. Notably, CAF4 and CAF12 were distinguished by their pronounced responsiveness to tumor-derived cues rather than by high basal invasive activity, suggesting the existence of distinct invasion-promoting and tumor-responsive fibroblast programs.

### Two fibroblast functional clusters are identified in the HNSCC TME

To obtain an integrated view of fibroblast functional heterogeneity, the analyzed features were combined to determine whether fibroblast populations segregate into distinct groups with preferential functional attributes. Plotting *contraction* against *invasion* identified two distinct CAF groups: highly invasive and contractile populations comprising CAF1, CAF2, CAF7, and CAF8, and low-invasive and low-contractile populations comprising CAF4 and CAF12. NFs were distributed between these extremes, representing intermediate phenotypic states **(Figure 3A)**. Fibroblast responsiveness to HNSCC-derived paracrine signals was then assessed by plotting the *stimulation index* against *invasion*. This analysis revealed that CAF4 and CAF12 again clustered together as highly responsive but low-invasive populations, whereas CAF2 and CAF8 displayed the opposite profile, characterized by high invasive capacity and limited responsiveness to tumor-derived factors **(Figure 3B).**

**Figure 3.**
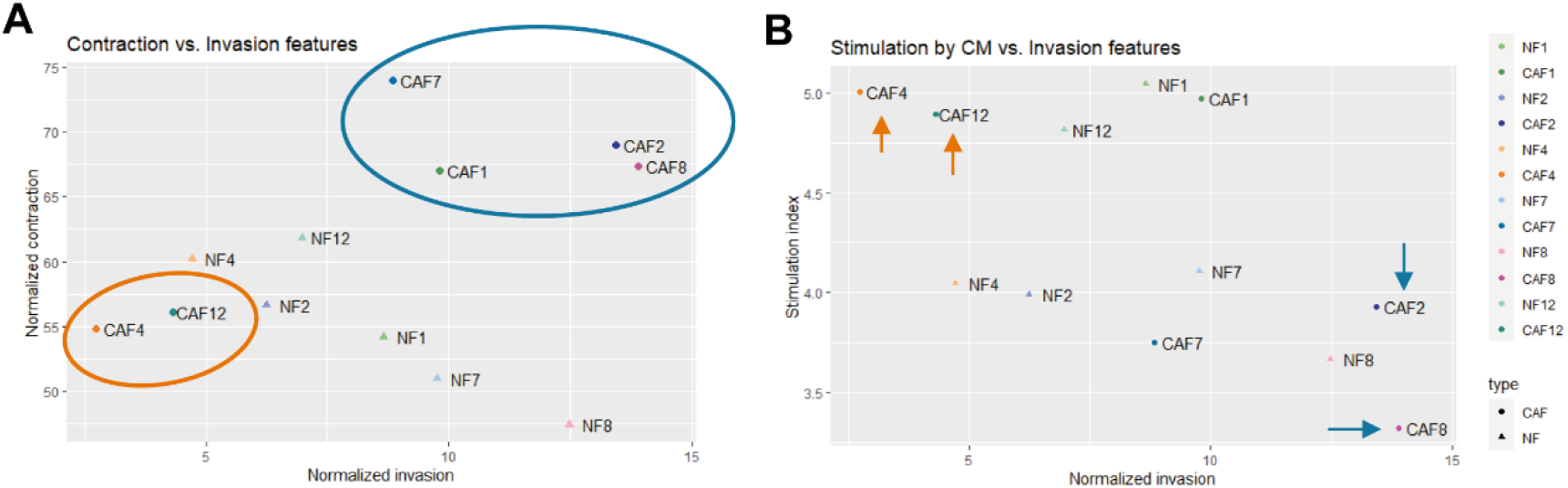
Fibroblast functional clusters. **A.** Scatter plot showing mean collagen contraction percentage *vs*. mean normalized invasion in each fibroblast population. **B.** Scatter plot showing stimulation index *vs*. mean normalized invasion in each fibroblast population. CAFs displaying polarized phenotypes are highlighted in orange and blue.

Integration of these analyses therefore defined two major CAF functional programs: an *Invasive* phenotype, characterized by high basal invasion and matrix-contractile activity but limited responsiveness to tumor-derived signals, representing an ECM-related phenotype, and a *Responsive* phenotype, characterized by low basal invasion and contractility but high responsiveness to tumor-derived paracrine signals. CAF2 and CAF8 were selected as representative of the *Invasive* phenotype, whereas CAF4 and CAF12 represented the *Responsive* phenotype. This functional classification was further supported at the molecular level, as Western blot analysis revealed recurrent and concordant protein-expression patterns that similarly distinguished CAF1, CAF2, CAF7, and CAF8 from the remaining populations **(Supplementary Figure S4)**, suggesting that protein marker expression levels align with their shared biological behavior. This supports the existence of common CAF-associated molecular programs underlying these functional groups.

### Proteomic profiling reveals candidate proteins associated with CAF-enriched ECM and adhesion programs

To characterize the molecular landscape underlying fibroblast heterogeneity, LC– MS/MS-based whole-proteome profiling was performed across all 12 primary fibroblast populations **(Supplementary Figure S5)**. Global differences between CAFs and NFs, patient-matched CAF-NF fibroblast pairs, and three additional comparisons guided by functional evidence derived from the *in vitro* assays were assessed via differential expression analysis (DEA). In the global CAF–NF comparison, 265 proteins were significantly differentially expressed, including 34 with |log_2_FC| > 1 **(Figure 4A)**. GO over-representation analysis revealed enrichment of processes related to cell adhesion, anchoring, locomotion, and ECM organization in CAFs, highlighting the predominance of adhesion- and matrix-associated programs **(Figure 4B).**

**Figure 4.**
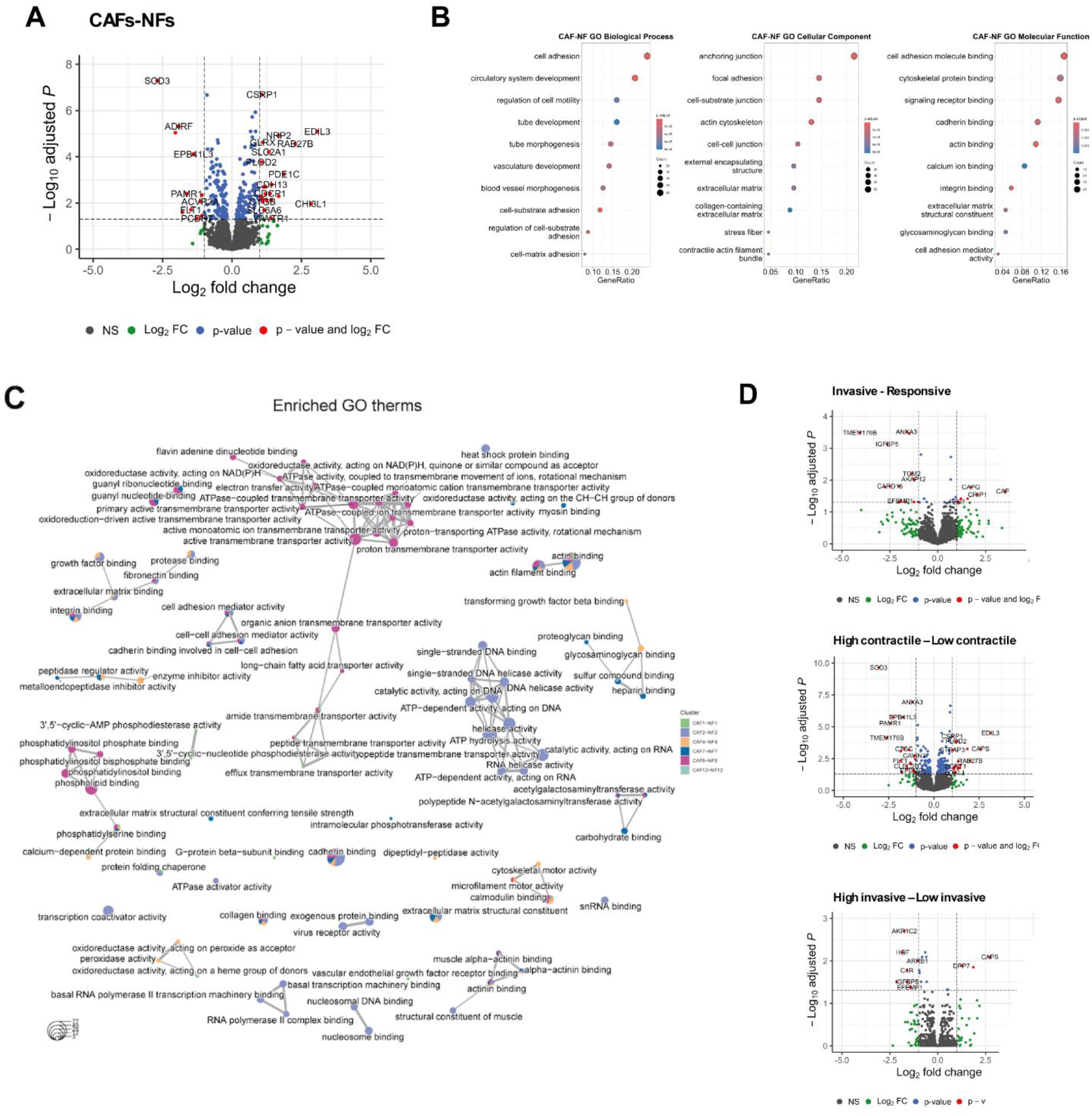
**Proteomic profiling of stromal fibroblasts**. **A**. Volcano plot showing differentially expressed (DE) proteins from all CAFs *vs*. all NFs comparison. Each dot represents a single protein. Grey dots indicate non-significant proteins; green dots indicate proteins with |log_2_FC| > 1; blue dots indicate proteins with adjusted *p*-value < 0.05; and red dots represent proteins with both |log_2_FC| > 1 and adjusted *p*-value < 0.05. **B.** Dot plots showing enriched terms from GO ORA using DE proteins with adjusted *p*-value < 0.05 as input, separated by each GO orthogonal ontology (i.e., molecular function (MF), biological process (BP), and cellular component (CC)). Top 10 most significant terms are displayed. Dot size represents gene counts and dot color represents *p*-values adjusted by Benjamini-Hochberg method. **C.** Emap plot of enriched GO terms using upregulated differentially expressed (DE) proteins in CAFs (adjusted *p*-value < 0.05 and log_2_FC > 0) from each paired comparison. After enrichment analysis, term similarity was calculated using the Jaccard correlation coefficient and represented as a network. Pie colors represent proportion of DE proteins from paired comparisons for a given term. **D.** Volcano plots showing differentially expressed (DE) proteins from Invasive *vs*. Responders, high *vs*. low Contractile and high *vs*. low Invasive comparisons. Figure keys as in (**A**).

Pairwise analyses further revealed substantial interpatient variability in the number of differentially expressed proteins between CAFs and their matched NFs, ranging from 114 to 1,052 across individual pairs. Of these, 57-376 proteins showed |log2FC| > 1. Consistent with the global CAF–NF comparison, processes related to cell adhesion, actin and integrin binding, ECM organization, and collagen-associated interactions were recurrently enriched across most paired comparisons **(Figure 4C)**.

To explore molecular features associated with the observed functional phenotypes, three additional comparisons were performed: C1, *Invasive* versus *Responsive*; C2, high versus low contractility; and C3, high versus low invasion **(Supplementary Figure S6)**. These analyses identified 34, 194, and 21 significantly differentially expressed proteins, respectively, of which 22, 38, and 12 showed |log2FC| > 1 **(Figure 4D)**.

Weighted gene co-expression network analysis further identified eight protein co-expression modules with distinct expression dynamics across fibroblast populations **(Figure 5A and B).** While some modules were strongly associated with individual populations, the green module showed a positive association across most CAFs, with the exception of CAF12 **(Figure 5C)**. Functional enrichment of this module again highlighted cell adhesion, anchoring, and actin-binding processes **(Figure 5D)**, reinforcing the recurrent association between CAF phenotypes and adhesion- and ECM-related molecular programs.

**Figure 5.**
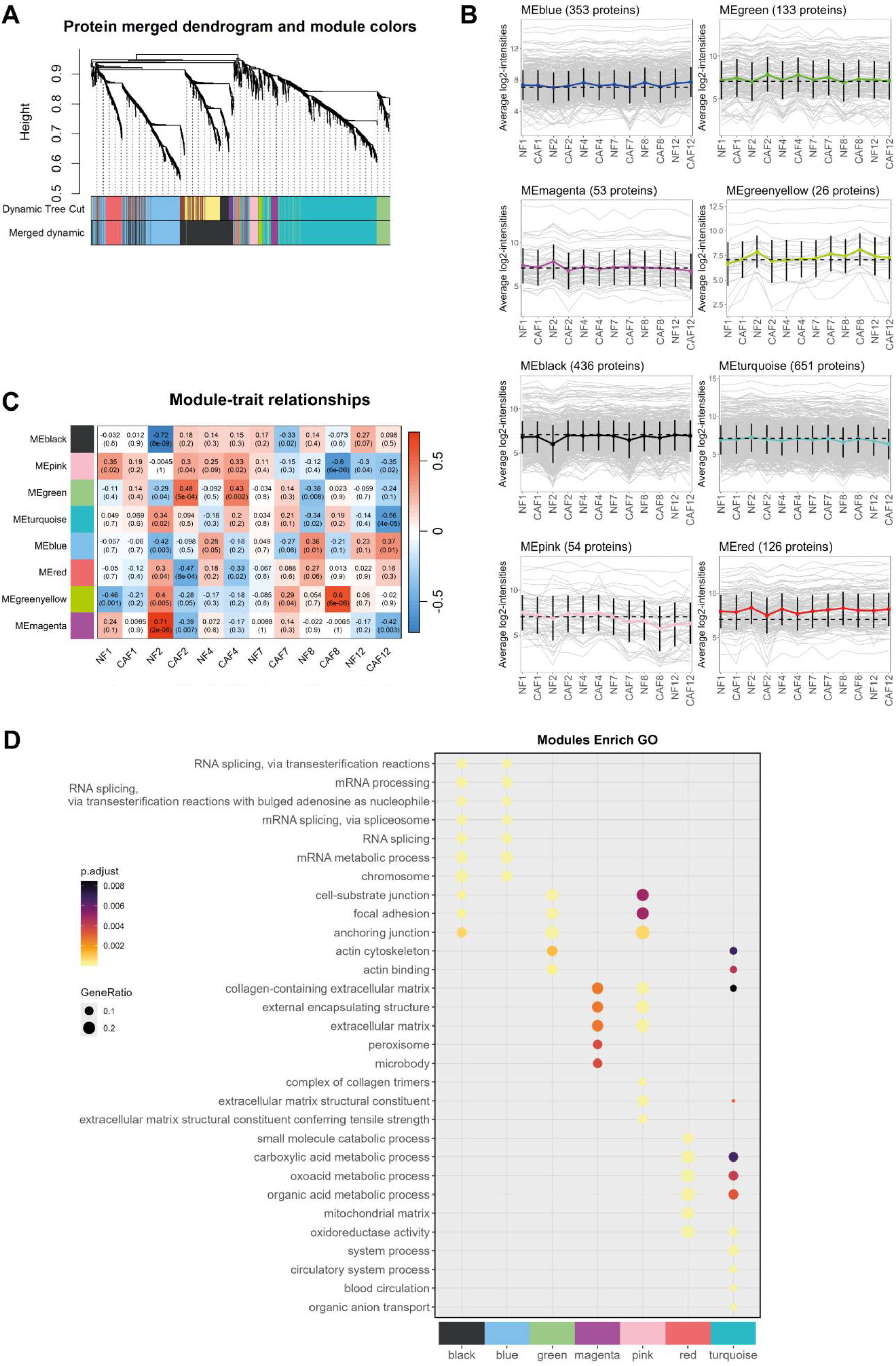
Protein co-expression network analysis. **A.** Dendrogram of network proteins and identified modules before (top) and after (bottom) cluster merging. Colors represent each different protein cluster (module). **B.** Dynamics of cluster expression profiles for each module. Each grey line represents a protein, and colored line represents the average of all proteins within the module. **C.** Module-trait relationship heatmap, showing correlations between protein modules and each fibroblast population. Positive and negative correlations are indicated in red and blue, respectively. Correlation coefficients and associated *p* values (in brackets) are indicated in each cell. **D.** Dot plot showing enriched terms from GO ORA using module proteins as input, separated by each co-expression cluster. Dot size represents gene ratio, and dot color represents Benjamini-Hochberg adjusted *p*-values.

### Pharmacological inhibition of protein candidates selectively modulates CAF-related pro-tumoral functions

Integration of proteomic and functional data across the twelve patient-matched primary fibroblast populations identified candidate proteins potentially associated with the observed phenotypic differences. We focused on pharmacologically targetable candidates to determine whether their inhibition could modulate CAF-related functional behavior **(Figure 6A)**.

**Figure 6.**
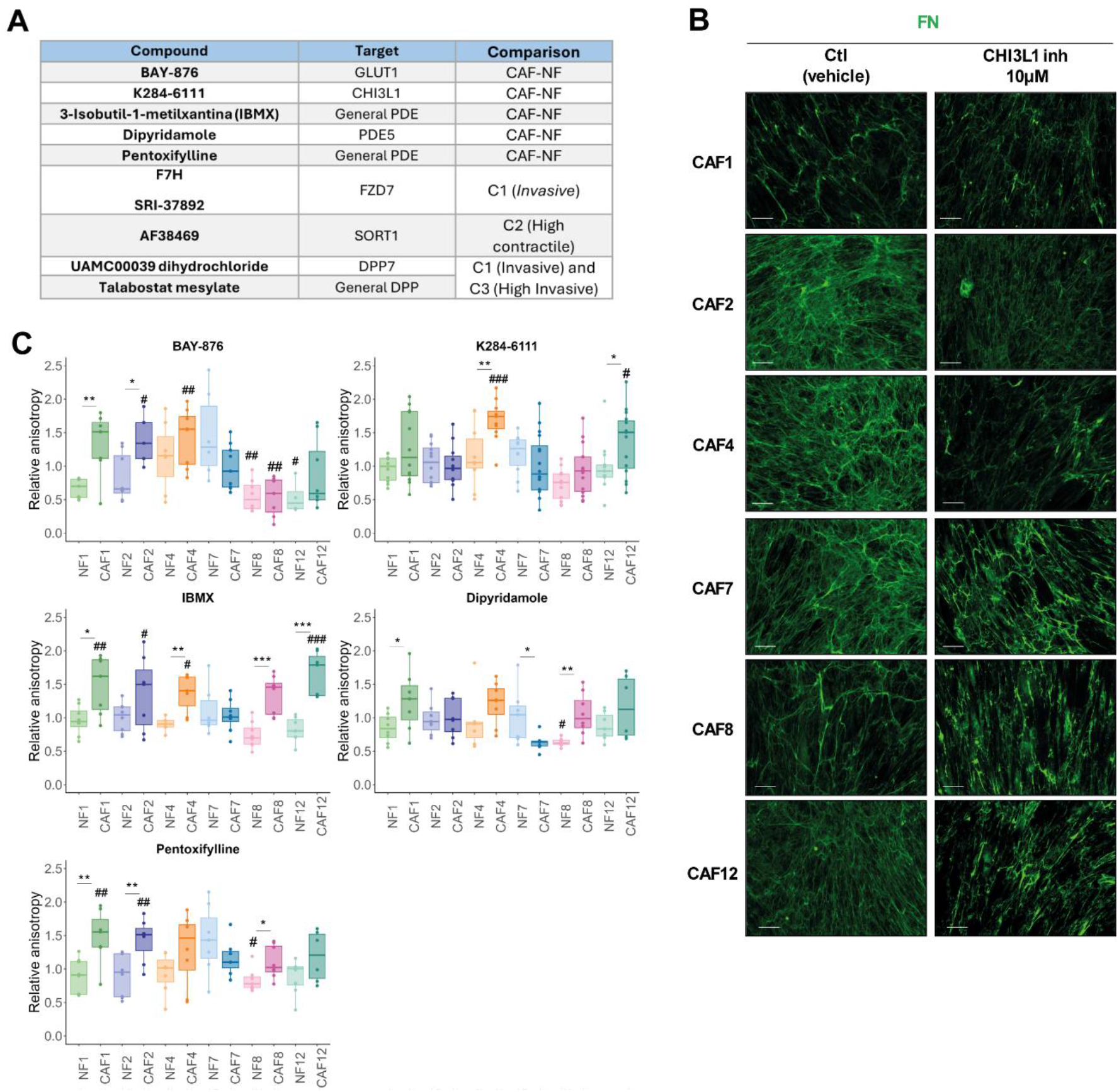
Effect of pharmacological inhibitors targeting CAF-related proteins on ECM architecture. **A**. Table displaying the selected protein targets, correspondent pharmacological inhibitors and the comparisons from proteomic data analysis where they emerged. **B.** Representative immunofluorescence images of fibronectin matrices derived from CAF populations after treatment with either vehicle (Ctl) or CHI3L1 inhibitor K284-6111 (10 µM). Green = fibronectin (FN). Scale bar = 50 µm. **C.** Boxplot showing relative anisotropy values for each fibroblast population after treatment with the following inhibitors: BAY-876, K284-6111, IBMX, dipyridamole and pentoxifylline. Anisotropy values were normalized to the corresponding control condition (vehicle) within each population; therefore, control values are not shown and correspond to a relative value of 1. Each dot represents a single image measurement. Measurements were performed on at least four images per assay in at least two independent experiments. Statistical significance indicated with # refers to comparisons between each treatment condition and the corresponding control, calculated using Dunnett’s multiple comparisons test. Statistical significance for CAF-NF comparisons is indicated with * and was calculated using unpaired *t*-test. * or # *p* < 0.05, ** or ## *p* < 0.01 and *** or ### *p* < 0.001.

We first evaluated targets emerging from the global CAF–NF comparison and examined the effects of GLUT1, CHI3L1, and PDE inhibition on ECM deposition and organization **(Supplementary Figure S7)**. Drug concentrations were selected to minimize cytotoxicity **(Supplementary Figure S8)**. Qualitative assessment of fibronectin matrices revealed a CAF-selective effect of the CHI3L1 inhibitor K284-6111, with little or no apparent effect on matched NFs. Marked alterations in ECM deposition and organization were observed in CAF2, CAF4, CAF7, CAF8, and CAF12, characterized by a less dense and more fragmented fibronectin network **(Figure 6B; Supplementary Figure S7)**.

Quantitative analysis of fiber anisotropy revealed distinct patterns of ECM modulation. Broad-spectrum PDE inhibition produced the most consistent effects on matrix organization, with IBMX increasing CAF-derived matrix anisotropy in all CAF populations except CAF7, while matched NFs remained largely unaffected. Pentoxifylline showed a similar but weaker pattern, whereas selective PDE5 inhibition with dipyridamole produced the least pronounced effects **(Figure 6C; Supplementary Figure S7)**. These findings indicate that matched CAF and NF populations can respond differently to the same pharmacological inhibition despite sharing the same patient genetic background and support a role for broad PDE signaling in CAF-mediated ECM organization. In contrast, GLUT1 inhibition with BAY-876 produced highly heterogeneous, population-dependent responses. CHI3L1 inhibition with K284-6111 significantly increased anisotropy in CAF4 and CAF12 **(Figure 6C; Supplementary Figure S7)**, however, these measurements did not fully capture the marked qualitative alterations in fibronectin deposition and architecture observed across several CAF populations. Because K284-6111 substantially reduced matrix deposition and disrupted the continuity of the fibronectin network, fiber-orientation measurements may be less informative under these conditions, as changes in matrix abundance and integrity can confound the assessment of fiber alignment.

Overall, these findings show that pharmacological inhibition can selectively modulate distinct aspects of CAF-derived ECM organization, with broad PDE inhibition increasing matrix alignment across several CAF populations and CHI3L1 inhibition primarily altering ECM deposition and architecture. However, these effects remained strongly population dependent; notably, CAF7 showed no consistent modulation of fiber anisotropy in response to any of the tested inhibitors. This variability highlights important differences in pharmacological sensitivity across patient-derived CAF populations and reinforces the need to account for interpatient heterogeneity when evaluating stromal-targeted strategies.

We next evaluated inhibitors targeting candidates derived from the functional group comparisons, including FZD7, SORT1, and DPP7 **(Supplementary Figure S6)**. Drug concentrations were selected to avoid major cytotoxic effects **(Supplementary Figure S9)**, and their impact was assessed on tumor-enhanced CAF invasion. Strikingly, FZD7 inhibition with either F7H or SRI-37892 completely abolished CAF spheroid invasion induced by Cal27-CM in all six primary CAF populations. F7H showed the strongest and most consistent inhibitory effect, reducing invasion below the levels observed in the non-stimulated condition in all CAF populations **(Figure 7)**. In contrast, SORT1 inhibition with AF38469 and DPP7 inhibition with UAMC00039 produced more modest and population-dependent effects, significantly reducing invasion in CAF4 and CAF7, and in CAF2 and CAF4, respectively **(Figure 7)**. These findings identify FZD7 inhibition as the most consistent pharmacological strategy for suppressing tumor-induced CAF invasion.

**Figure 7.**
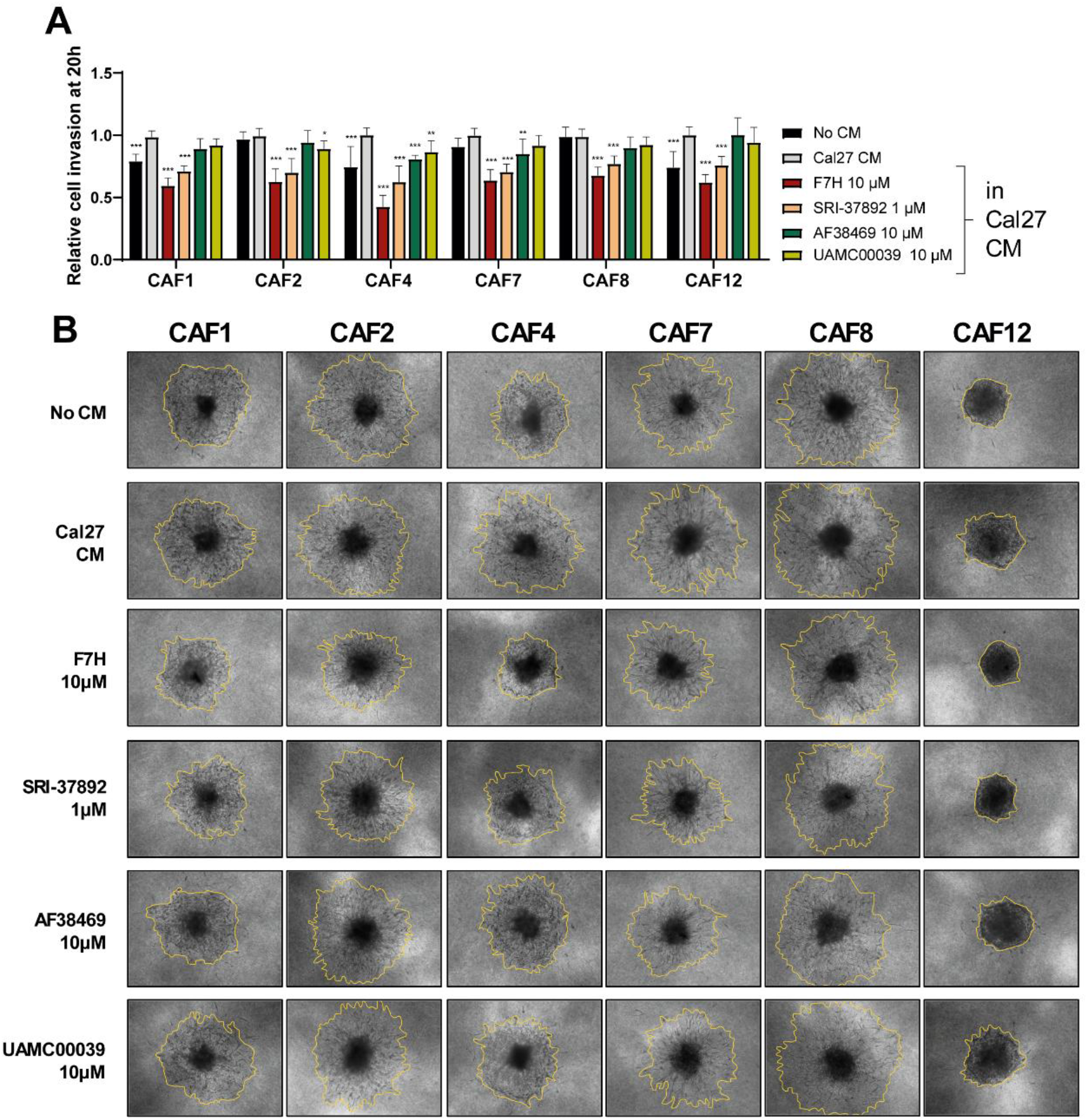
Effect of pharmacological inhibitors targeting functional candidates on HNSCC-driven CAF invasion. **A.** 3D spheroid invasion of primary CAF populations was monitored for 20 h in the absence (No CM) or presence of Cal27 CM, untreated or treated with the indicated inhibitors (1 µM or 10 µM). Bar chart showing CAF spheroid invasion at 20 h, normalized to area at 0 h and expressed relative to the corresponding Cal27 CM (untreated condition). Data are represented as mean ± SD. Statistical significance for each treatment condition compared to Cal27 CM condition was calculated using Dunnett’s multiple comparisons test. \**p* < 0.05, ** *p* < 0.01 and \*\*\**p* < 0.001. n = 3 independent experiments in triplicate. **B.** End-point representative images of CAF spheroids at 20 h for the different conditions tested. The final CAF invading area is highlighted in yellow.

## DISCUSSION

In this study, we investigated functional and molecular CAF heterogeneity in HNSCC using patient-matched primary CAF and NF populations, enabling direct within-patient comparisons while capturing interpatient variability across independent fibroblast populations.

Our data revealed marked functional diversity among fibroblast populations. Some displayed high basal invasive and contractile activity, whereas others showed lower basal activity but pronounced responsiveness to tumor-derived cues. NFs also exhibited interpatient heterogeneity and variable plasticity in response to tumor signals, consistent with previous reports showing that normal or peritumoral fibroblasts can acquire tumor-promoting functions upon activation (29,30).

Among the analyzed traits, ECM remodeling emerged as one of the most consistent CAF-associated features. CAFs generated denser and more disorganized fibronectin matrices than matched NFs and induced less compact, more interspersed tumor nest architectures. These findings agree with previous studies linking increased ECM deposition, stiffness, fibronectin organization, and collagen remodeling with tumor progression, CAF activation, immune exclusion, and poor prognosis (31–36). Proteomic analyses further supported this phenotype, revealing recurrent enrichment of adhesion-, cytoskeleton-, and ECM-related programs in CAFs.

Integration of the functional assays identified two major CAF phenotypes. The *Invasive* group, represented by CAF2 and CAF8, combined high basal invasion and contractility, whereas the *Responsive* group, represented by CAF4 and CAF12, showed lower basal activity but marked responsiveness to tumor-derived factors. Similar functional diversification has been described in oral cancer CAF subsets with distinct migratory and secretory properties (30), supporting the existence of multiple tumor-promoting CAF phenotypes.

Rather than approaching CAF heterogeneity exclusively from a molecular perspective, we integrated functional phenotypes with proteomic data to identify pharmacologically targetable candidates associated with specific CAF behaviors. Given the increasing interest in CAF targeting as a complementary anticancer strategy (37–45), we evaluated whether inhibition of selected candidates could modulate distinct CAF-mediated pro-tumoral functions.

CHI3L1 inhibition markedly altered fibronectin deposition and matrix architecture, with particularly pronounced effects in CAF7 and CAF8, both displaying strong ECM-related features. CHI3L1 has previously been associated with fibrosis and pro-tumoral CAF activity in several cancers (46–49), and our findings extend its potential role to ECM remodeling in HNSCC-derived CAFs. Broad PDE inhibition increased matrix anisotropy across several CAF populations while having limited effects on matched NFs. Previous studies have linked PDE inhibition to reduced myofibroblast activation, contractility, and ECM production in fibrotic diseases and cancer (50–53). Together, these findings show that distinct aspects of CAF-mediated ECM remodeling can be pharmacologically modulated, while the variable responses across patient-derived populations highlight substantial heterogeneity in stromal drug sensitivity.

FZD7 emerged as the most consistent pharmacological vulnerability identified in our functional analyses. Inhibition with either F7H or SRI-37892 robustly abrogated tumor-induced CAF invasion across all tested populations. Wnt signaling has been implicated in fibrosis and CAF activation (34,54–56), but, to our knowledge, this is the first study linking FZD7 to tumor-induced CAF invasion in HNSCC. Given the established contribution of CAF invasion to tumor dissemination (30,57,58), these findings support FZD7 as a promising target for suppressing pro-invasive stromal behavior.

Further studies using single-cell approaches and more complex tumor–stroma models will help resolve intra-patient fibroblast diversity and validate these functional programs in increasingly physiologically relevant contexts. Nevertheless, our patient-derived models reproducibly captured functional and molecular CAF heterogeneity and enabled the identification of pharmacologically targetable vulnerabilities.

## CONCLUSIONS

Taken together, our findings demonstrate that HNSCC-derived CAFs display marked functional and molecular heterogeneity, with ECM remodeling emerging as a recurrent CAF-associated feature. Integration of functional and proteomic profiling identified distinct CAF programs and pharmacologically targetable vulnerabilities capable of modulating specific pro-tumoral functions. While some responses were strongly population dependent, FZD7 inhibition consistently suppressed tumor-induced CAF invasion. These findings support functional stratification of CAFs as a framework for developing stromal-targeted strategies and future combination approaches for precision oncology directed against both tumor cells and pro-tumoral stromal programs.

## Supporting information

Supplementary_information_Prieto_Fernandez

## Abbreviations

ACN: acetonitrile
BSA: bovine serum albumin
CAF: cancer-associated fibroblast
CHI3L1: chitinase 3-like 1
CM: conditioned media
DAPI: 4′,6-diamidino-2-phenylindole
DEA: differential expression analysis
DPP7: dipeptidyl peptidase 7
DTT: dithiothreitol
ECM: extracellular matrix
EGFR: epidermal growth factor receptor
FBS: fetal bovine serum
FDA: Food and Drug Administration
FFPE: formalin-fixed paraffin-embedded
FN: fibronectin
FZD7: Frizzled Class Receptor 7
GO: Gene Ontology
HNSCC: head and neck squamous cell carcinoma
HPV: human papillomavirus
ITS: insulin-transferrin-selenium
LC-MS/MS: liquid chromatography–tandem mass spectrometry
LSCC: laryngeal squamous cell carcinoma
MTS: 3-(4,5-dimethylthiazol-2-yl)-5-(3-carboxymethoxyphenyl)-2-(4-sulfophenyl)-2H-tetrazolium
NF: normal fibroblast
ORA: over-representation analysis
OS: overall survival
PCA: principal component analysis
PDE: phosphodiesterase
PDE5: phosphodiesterase 5
RT: room temperature
SORT1: sortilin 1
TFA: trifluoroacetic acid
TME: tumor microenvironment
WGCNA: weighted gene co-expression network analysis.

## DECLARATIONS

### Ethics approval and consent to participate

All human samples used in this study were obtained through the Principado de Asturias Biobank (BioPA, National Registry of Biobanks B.0000827; PT20/00161 and PT23/00077). Written informed consent was obtained from all patients. The study was conducted in accordance with the principles of the Declaration of Helsinki and approved by the Ethics Committee of the Hospital Universitario Central de Asturias and by the Regional CEIC from Principado de Asturias (CEImPA; approval numbers 2023.018 for project PI22/00167 and 2024.187 for project PI24/00398). All patient information was handled confidentially in accordance with institutional and biobank guidelines.

### Data and Code Availability Statement

The mass spectrometry proteomics data generated in this study have been deposited to the ProteomeXchange Consortium via the PRIDE partner repository with the dataset identifier PXD078734. The dataset is currently private and will be made publicly available upon publication of the manuscript. The code used for proteomics data analysis is available at: https://github.com/llarapf/NF-CAF-characterization-proteomics, currently in private status, access is available upon requested token. Additional data supporting the findings of this study are available from the corresponding author upon reasonable request, subject to privacy and ethical restrictions.

### Authors’ contributions

L.P.-F.: Conceptualization, Methodology, Investigation, Formal analysis, Data curation, Visualization, Writing – original draft, Writing – review & editing. A.M.-C.: Investigation, Resources, Writing – review & editing. L.V.: Investigation, Resources, Writing – review & editing. A.G.-T.: Methodology, Investigation, Formal analysis, Writing – review & editing. B.L.-D.: Methodology, Investigation, Formal analysis, Writing – review & editing. F.H.-P.: Formal analysis, Writing – review & editing. I.N.-L.: Methodology, Resources, Writing – review & editing. C.R.: Methodology, Resources, Writing – review & editing. R.G.-E.: Resources, Supervision, Writing – review & editing. J.P.R.: Resources, Supervision, Funding acquisition, Writing – review & editing. J.C.V.: Investigation, Resources, Supervision, Funding acquisition, Writing – review & editing. T.R.-S.: Investigation, Resources, Supervision, Writing – review & editing. J.M.G.-P.: Conceptualization, Supervision, Funding acquisition, Writing – original draft, Writing – review & editing. S.A.-T.: Conceptualization, Methodology, Investigation, Formal analysis, Project administration, Supervision, Funding acquisition, Writing – original draft, Writing – review & editing.

## Acknowledgements

The authors thank the patients and their families for their participation in this study. We also thank Dr. Reidar Grenman (University of Turku, Finland) for kindly providing the UT-SCC40 cell line. We particularly acknowledge the collaboration of the Principado de Asturias Biobank (PT20/00161 and PT23/00077), part of the Spanish National Biobanks Network, jointly financed by the Servicio de Salud del Principado de Asturias, Instituto de Salud Carlos III and Fundación Bancaria Cajastur.

## Additional files

**Supplementary Information.** Supplementary Tables S1-S4. Supplementary Figures S1-S9.

