## Supplementary_information_Prieto_Fernandez for "Functional proteomics identifies targetable cancer-associated fibroblast programs in head and neck cancer"

Prieto-Fernández *et al.*

**The PDF file includes:**

Tables S1, S2, S3 and S4

Figs. S1, S2, S3, S4, S5, S6, S7, S8, and S9

**Table S1. Clinicopathological characteristics of HNSCC patients.**

| PATIENT NUMBER | TISSUE ORIGIN | TNM | STAGE | HPV STATUS | SEX | AGE |
|---|---|---|---|---|---|---|
| **1** | Oral cavity | T4N1M0 | IV A | Negative | Male | 53 |
| **2** | Oral cavity | T3N2bM0 | IV A | Negative | Male | 66 |
| **4** | Oral cavity | T3N3bM0 | IV B | Negative | Male | 65 |
| **7** | Oral cavity | T4N0M0 | IV A | Negative | Male | 61 |
| **8** | Oral cavity | T3N2bM0 | IV A | Negative | Male | 87 |
| **12** | Oral cavity | T3N3bM0 | IV B | Negative | Female | 68 |

**Table S2. Primary and secondary antibodies used for immunofluorescence**

| Antibody | Supplier (Reference) | Host species | Working concentration |
|---|---|---|---|
| **Primary antibodies** | | | |
| **Fibronectin** | Abcam (ab2413) | Rabbit | 1:200 |
| **Vimentin** | Abcam [RV202] (ab8978) | Mouse | 1:200 |
| **Secondary antibodies** | | | |
| **Alexa Fluor® 488 goat anti-rabbit** | Thermo Fisher (A11008) | Goat | 1:200 |
| **Alexa Fluor® 555 goat anti-mouse** | Thermo Fisher (A21422) | Goat | 1:200 |

**Table S3. Pharmacological inhibitors used for *in vitro* assays**

| Compound | Supplier (Reference) | Target |
|---|---|---|
| BAY-876 | MedChemExpress (HY-100017) | GLUT1 |
| K284-6111 | MedChemExpress (HY-148013) | CHI3L1 |
| 3-Isobutil-1-metilxantina (IBMX) | MedChemExpress (HY-12318) | General PDE |
| Dipyridamole | Sigma (D9766) | PDE V |
| Pentoxifylline | Sigma (P1784) | General PDE |
| F7H | MedChemExpress (HY-156095) | FZD7 antagonist, extracellular binding |
| SRI-37892 | MedChemExpress (HY-117002) | FZD7 inhibitor, Wnt-FZD7 interaction |
| AF38469 | MedChemExpress (HY-12802) | SORT1 |
| UAMC00039 dihydrochloride | MedChemExpress (HY-101769) | DPP7 |

**Table S4. Primary and secondary antibodies used for Western Blot analysis**

| Antibody | Supplier (Cat. No.) | Molecular weight (kDa) | Host species | Working Concentration |
|---|---|---|---|---|
| **Primary antibodies** | | | | |
| **Fibronectin** | Abcam (ab2413) | 262 | Rabbit | 1:1,000 |
| **PDGFRα** | Abcam (ab203491) | 150 | Rabbit | 1:1,000 |
| **Integrinβ1 (CD29)** | Abcam (ab52971) | 120-140 | Rabbit | 1:1,000 |
| **LOXL2** | Thermofisher (PA5-85210) | 100-120 | Rabbit | 1:1,000 |
| **FAP** | Abcam (ab207178) | 100 | Rabbit | 1:1,000 |
| **N-cadherin** | Novus Biological (NCNBP1-48309) | 100 | Mouse | 1:1,000 |
| **β-catenin** | BD Biosciences (610153) | 95 | Mouse | 1:1,000 |
| **YAP** | Invitrogen (PA1-46189) | 70 | Rabbit | 1:1,000 |
| **Vimentin** | Abcam [RV202] (ab8978) | 57 | Mouse | 1:1,000 |
| **α-SMA** | Sigma (A2547) | 42 | Mouse | 1:1,000 |
| **GAPDH clone 6C5** | Merck Millipore (MAB374) | 36 | Mouse | 1:1,000 |
| **pMLC2** | Cell Signaling (3671) | 20 | Rabbit | 1:1,000 |
| **SDF-1** | Abcam (ab9797) | 15 | Rabbit | 1:1,000 |
| **S100A4** | Abcam (ab124805) | 12 | Rabbit | 1:1,000 |
| **Secondary antibodies** | | | | |
| **IRDye® 680RD Goat Anti-Mouse IgG** | LICORbio (926-68070) | - | Goat | 1:10,000 |
| **IRDye® 800CW Goat Anti-Mouse IgG** | LICORbio (926-32210) | - | Goat | 1:10,000 |
| **IRDye® 800CW Goat Anti-Rabbit IgG** | LICORbio (926-32211) | - | Goat | 1:10,000 |

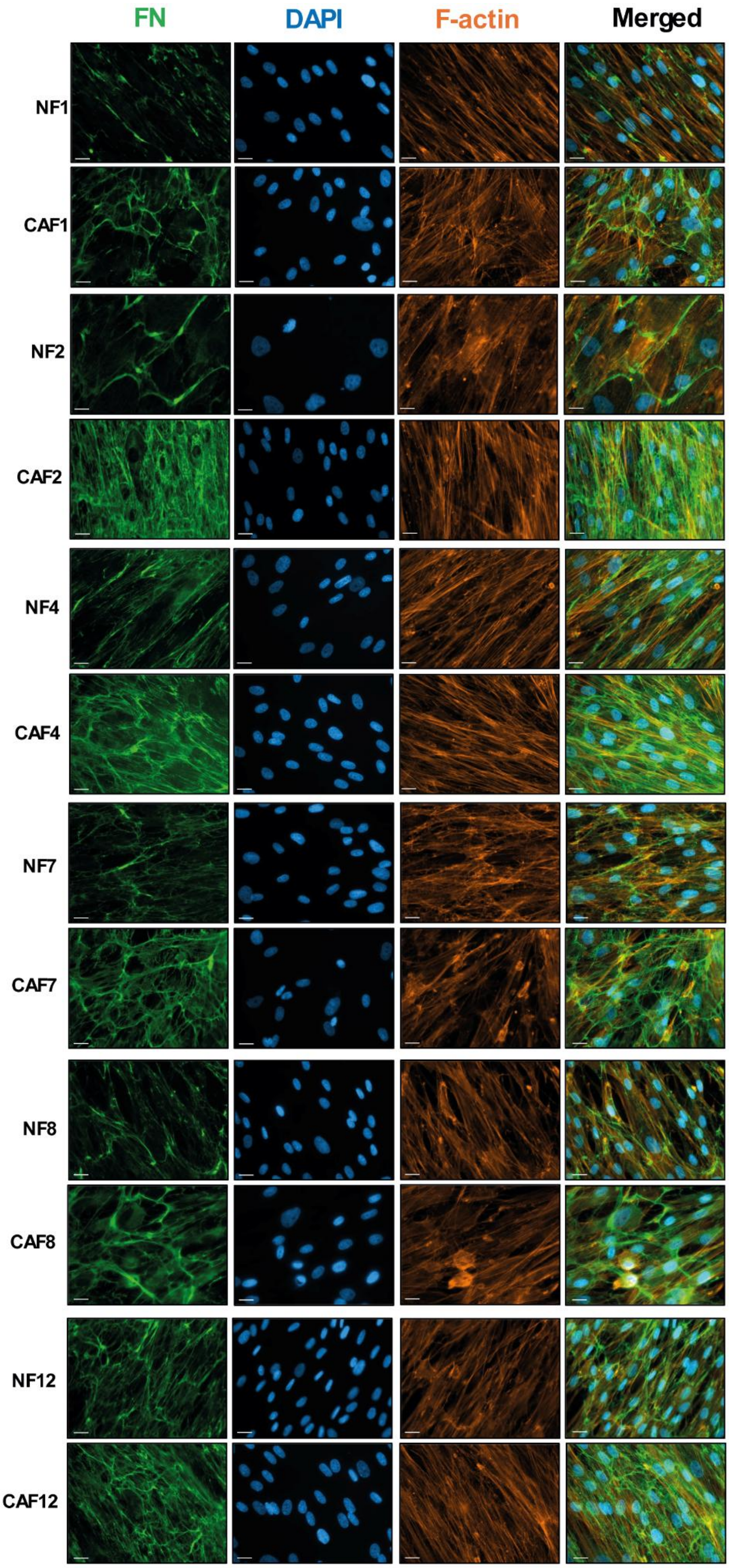
FN
DAPI
F-actin
Merged
NF1
CAF1
NF2
CAF2
NF4
CAF4
NF7
CAF7
NF8
CAF8
NF12
CAF12

**Supplementary Figure S1. Fibronectin ECM synthesis and deposition by NFs and CAFs.** Representative immunofluorescence images of fibroblast-derived fibronectin ECM after 5 days in culture. n = 3 independent experiments. Green = fibronectin (FN); blue = DAPI; orange = F-actin. Scale bar = 20 μm.

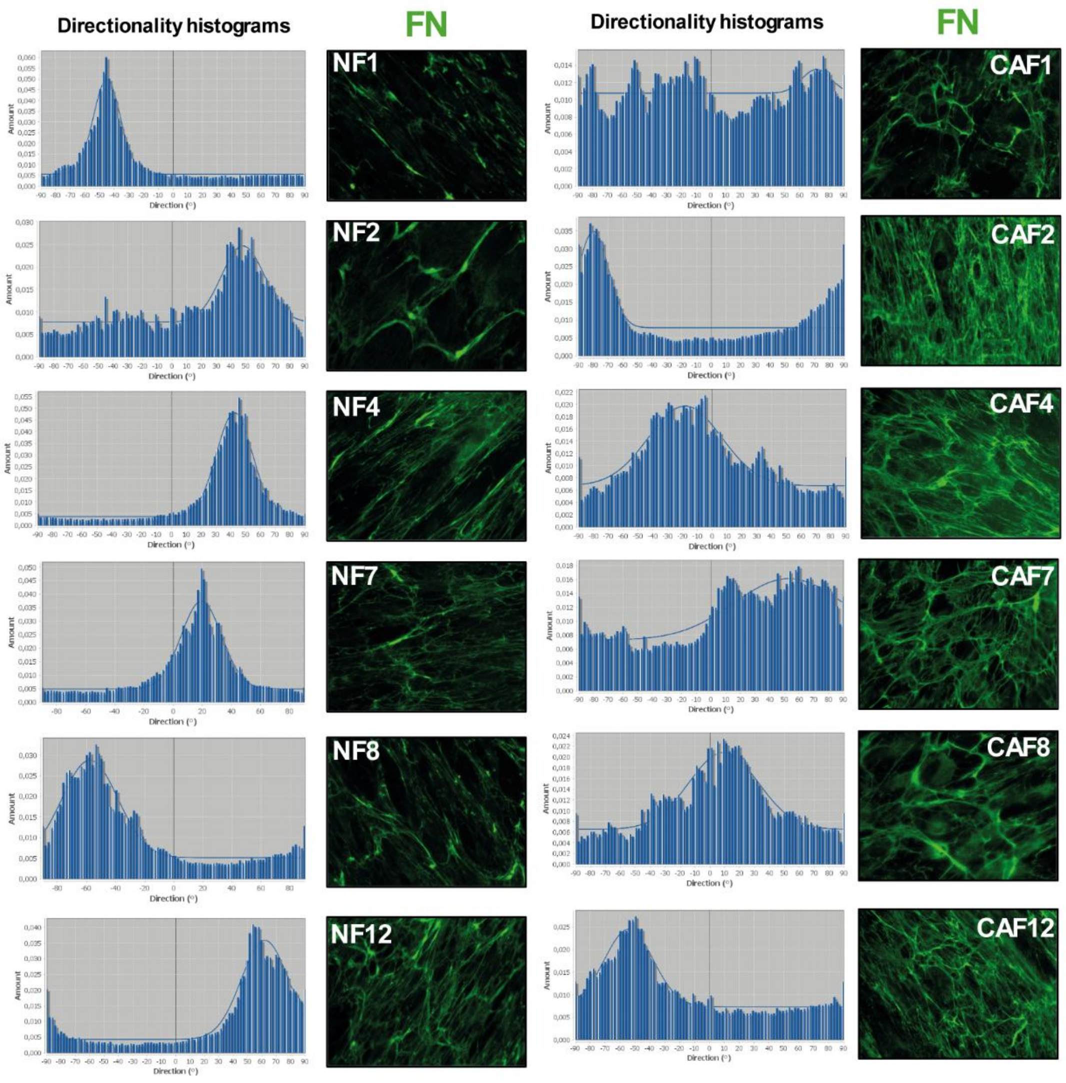


**Supplementary Figure S2. Directionality of fibronectin fibers in fibroblast-derived matrices.** Representative immunofluorescence images of fibronectin staining (green) in NF- and CAF-derived matrices, together with the corresponding fiber directionality histograms generated using the ImageJ Directionality plugin.

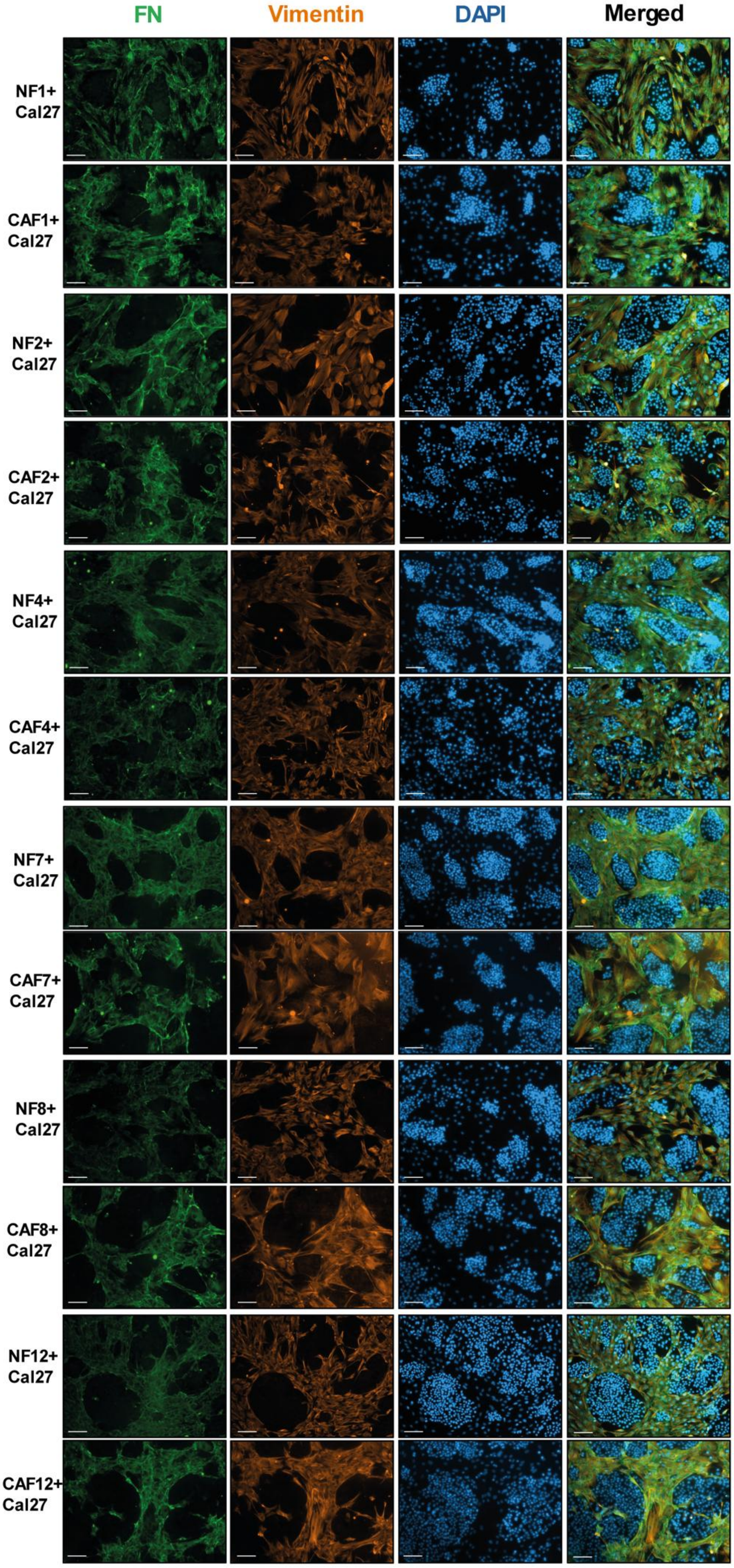

FN
Vimentin
DAPI
Merged
NF1+ Cal27
CAF1+ Cal27
NF2+ Cal27
CAF2+ Cal27
NF4+ Cal27
CAF4+ Cal27
NF7+ Cal27
CAF7+ Cal27
NF8+ Cal27
CAF8+ Cal27
NF12+ Cal27
CAF12+ Cal27

**Supplementary Figure S3. Tumor nest architecture in HNSCC cell–fibroblast co-cultures.** Representative immunofluorescence images of co-cultured Cal27 cells with the indicated fibroblast population (proportion 1:1) after 72 h. Green = fibronectin (FN); blue = DAPI; orange= Vimentin. Scale bar = 100 μm.

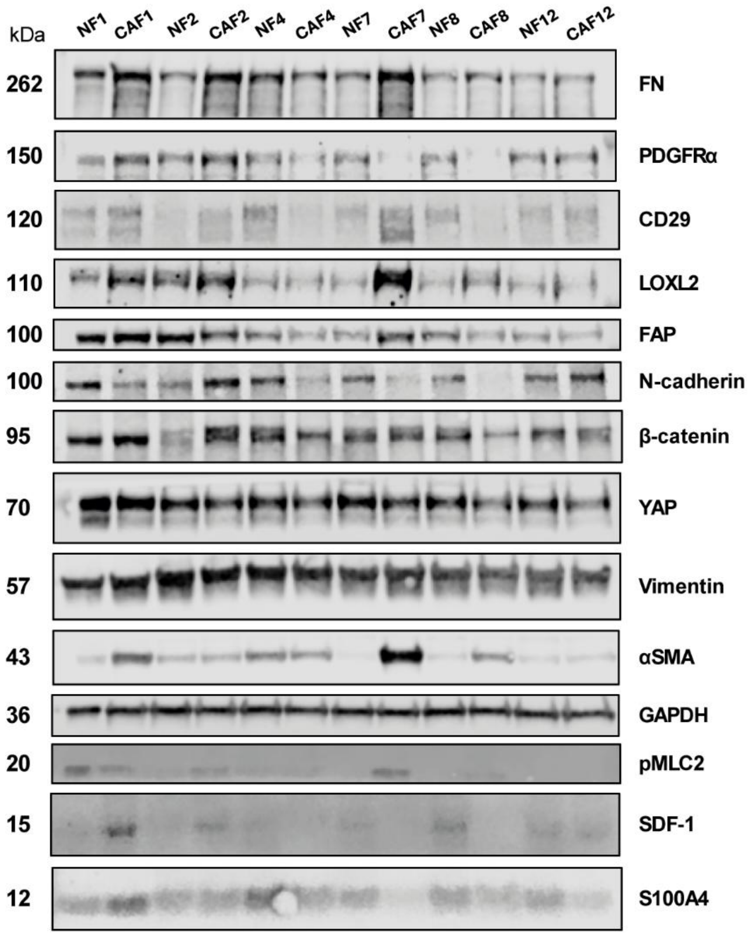


**Supplementary Figure S4. Analysis of CAF-related markers in patient-matched primary fibroblast populations**. Protein expression of indicated CAF markers was assessed by Western Blot. GAPDH was used as loading control. Molecular weights for each protein are displayed on the left in kilodaltons (kDa).

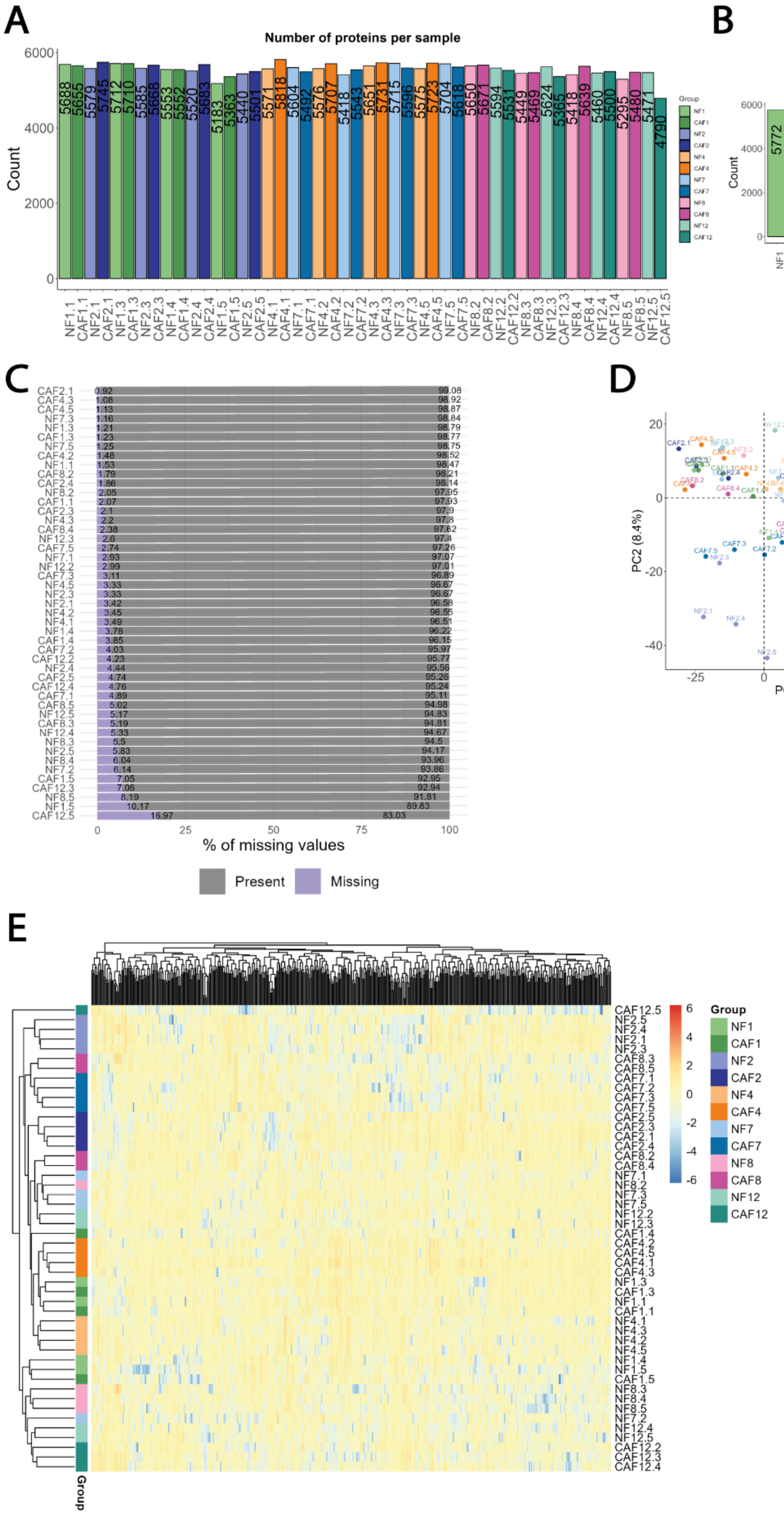

A
Number of proteins per sample
Count
6000
4000
2000
0
5688 5655 5579 5745 5712 5710 5585 5668 5553 5552 5520 5683 5183 5363 5440 5501 5571 5818 5604 5492 5576 5707 5418 5543 5651 5731 5715 5596 5575 5723 5704 5618 5650 5671 5594 5531 5449 5469 5624 5365 5418 5639 5460 5500 5295 5480 5471 4790
NF1.1 CAF1.1 NF2.1 CAF2.1 NF1.3 CAF1.3 NF2.3 CAF2.3 NF1.4 CAF1.4 NF2.4 CAF2.4 NF1.5 CAF1.5 NF2.5 CAF2.5 NF4.1 CAF4.1 NF7.1 CAF7.1 NF4.2 CAF4.2 NF7.2 CAF7.2 NF4.3 CAF4.3 NF7.3 CAF7.3 NF4.5 CAF4.5 NF7.5 CAF7.5 NF8.2 CAF8.2 NF12.2 CAF12.2 NF8.3 CAF8.3 NF12.3 CAF12.3 NF8.4 CAF8.4 NF12.4 CAF12.4 NF8.5 CAF8.5 NF12.5 CAF12.5
Group
NF1 CAF1 NF2 CAF2 NF4 CAF4 NF7 CAF7 NF8 CAF8 NF12 CAF12
B
Number of proteins per Group
Count
6000
4000
2000
0
5772 5781 5746 5794 5763 5818 5783 5752 5746 5757 5749 5722
NF1 CAF1 NF2 CAF2 NF4 CAF4 NF7 CAF7 NF8 CAF8 NF12 CAF12
C
% of missing values
0 25 50 75 100
Present
Missing
D
PCA
PC1 (19.4%)
PC2 (8.4%)
E
Group
NF1 CAF1 NF2 CAF2 NF4 CAF4 NF7 CAF7 NF8 CAF8 NF12 CAF12
6 4 2 0 -2 -4 -6

**Supplementary Figure S5. LC-MS/MS proteomics quality control and dimensionality reduction of fibroblast proteomes**. **A.** Bar plot representing the number of proteins identified per sample. **B.** Bar plot showing the number of proteins identified per group. **C**. Representation of missing value percentages in each sample. **D.** Principal component analysis (PCA) representation of whole imputed protein expression matrix after pre-processing steps. **E**. Heatmap representing column-scaled expression of the 500 proteins with highest variance across samples. Similarity between columns and rows was calculated with Euclidean distances and complete-linkage was used for hierarchical clustering.

| | | | | | |
|---|---|---|---|---|---|
| **C1** | Invasive (I) | CAF2, CAF8 | *vs.* | Responsive (R) | CAF4, CAF12 |
| **C2** | High contractile (hC) | CAF1, CAF2, CAF7, CAF8 | | Low contractile (lC) | NF1, NF2, NF7, NF8, CAF4, CAF12 |
| **C3** | High invasive (hI) | CAF2, CAF8, NF8 | | Low invasive (lI) | NF2, NF4, NF12, CAF4, CAF12 |

**Supplementary Figure S6. Comparisons for differential expression analysis modelling *in vitro* functional evidence**. Table displaying the comparisons for differential expression analysis modeling *in vitro* functional evidence, with the fibroblast populations belonging to each group indicated.

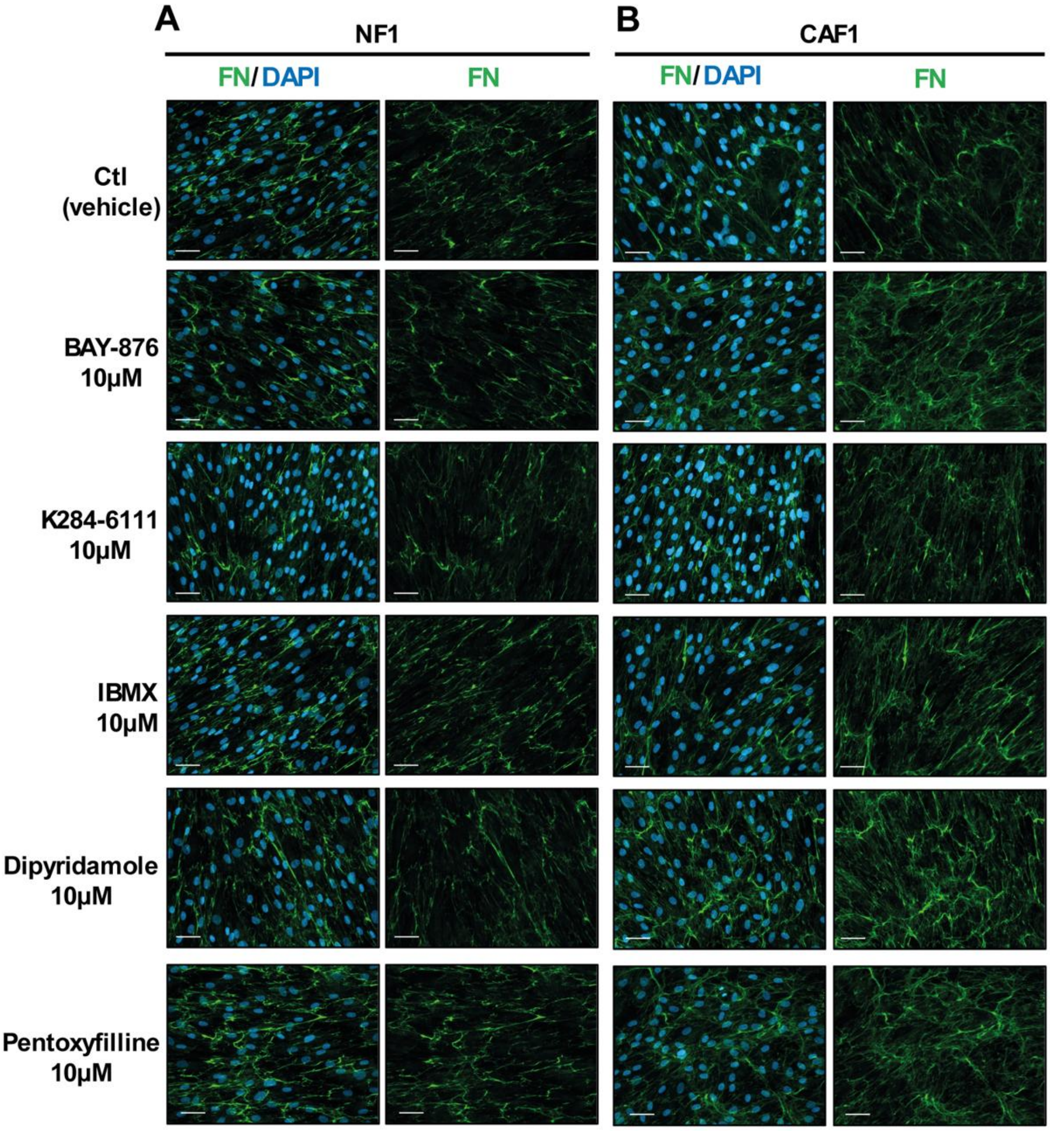


Figure continues next page ▶

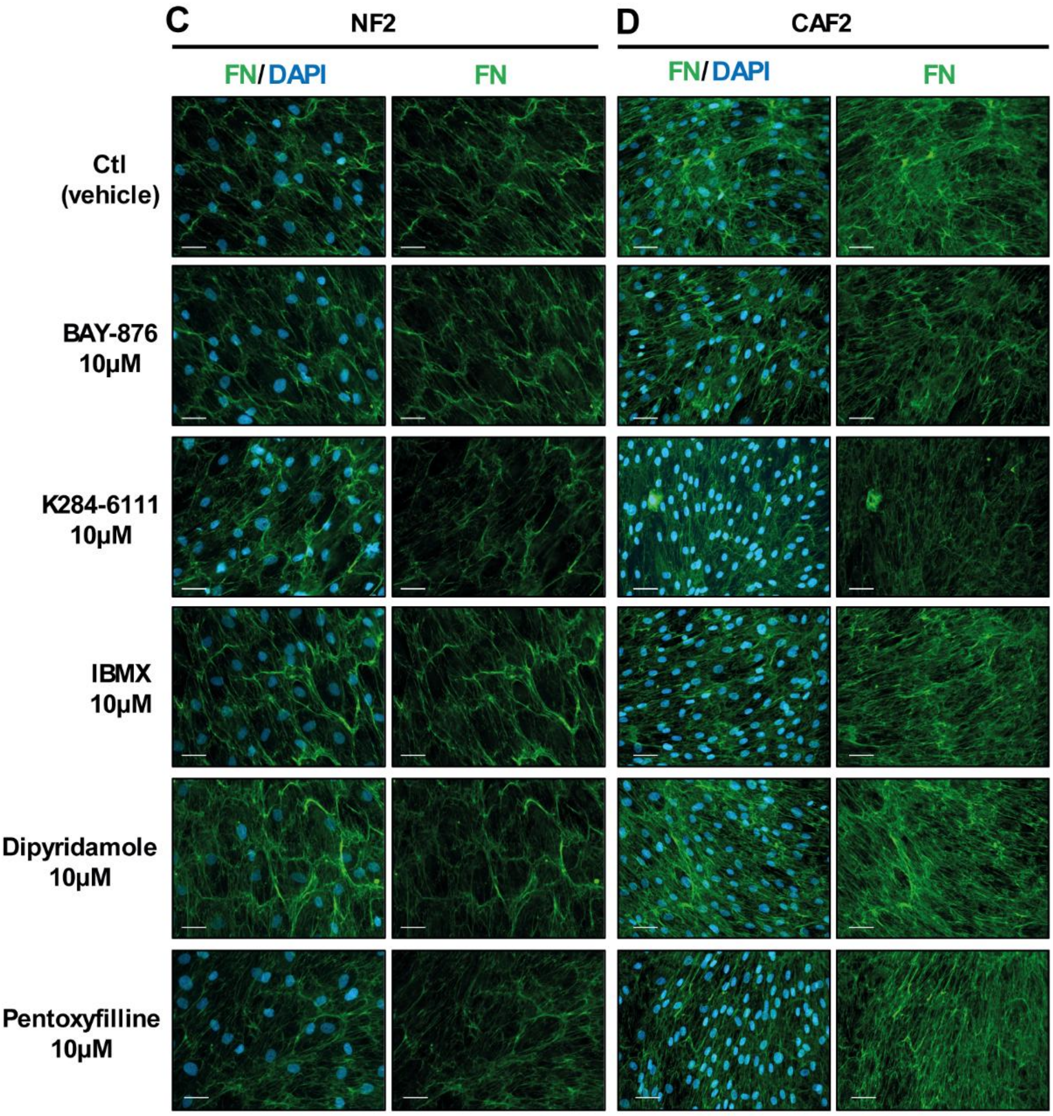


**Figure continues next page** ▶

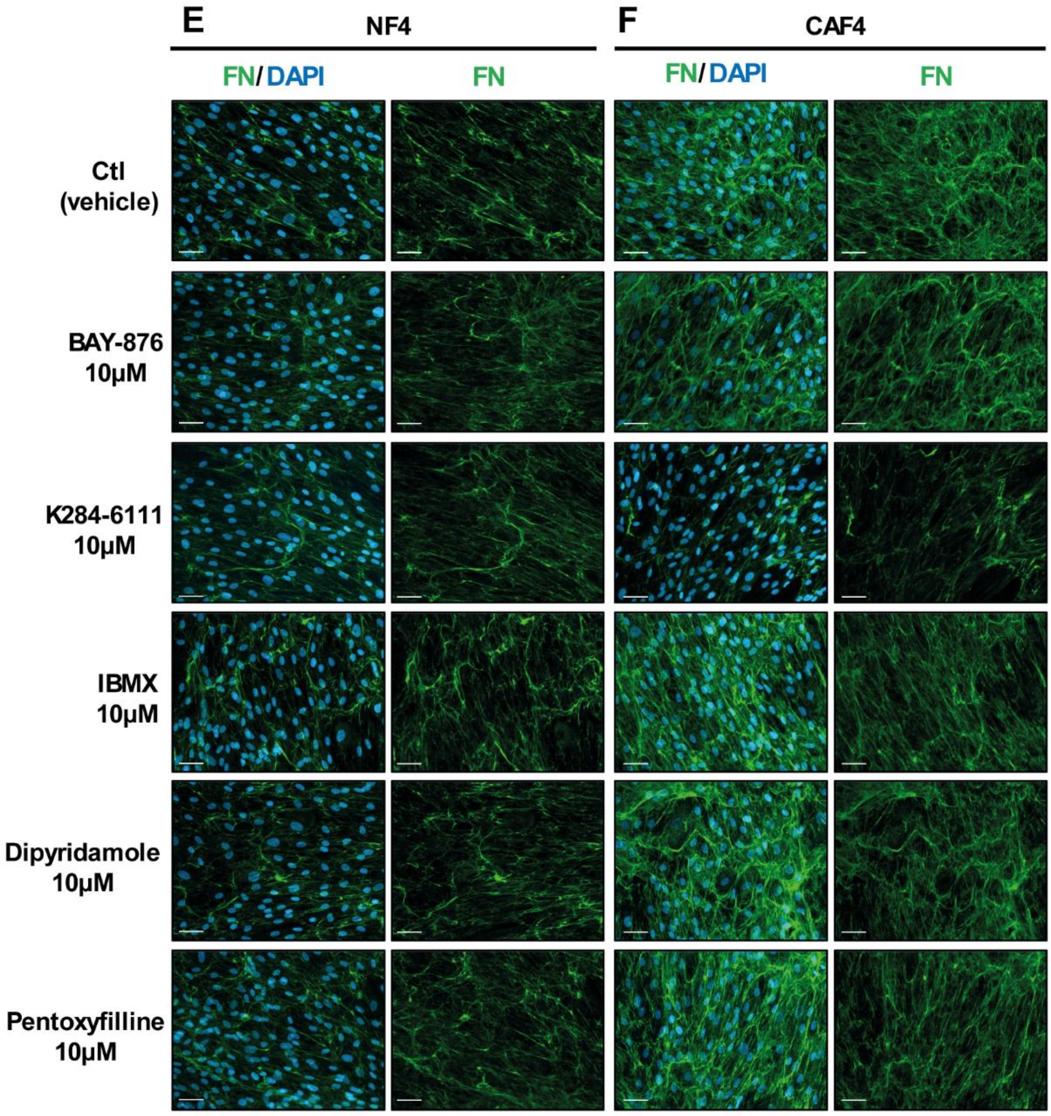


Figure continues next page ▶

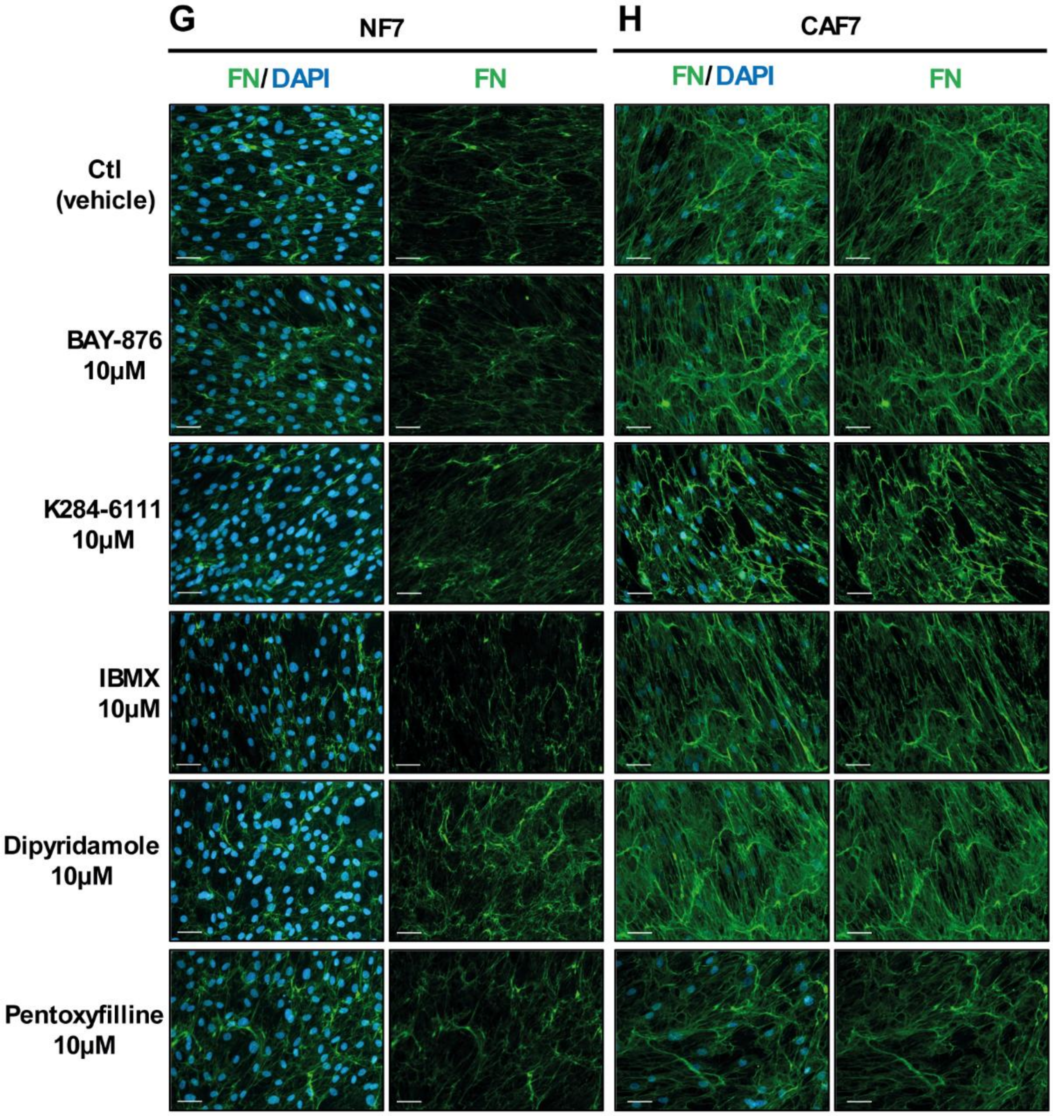


Figure continues next page ▶

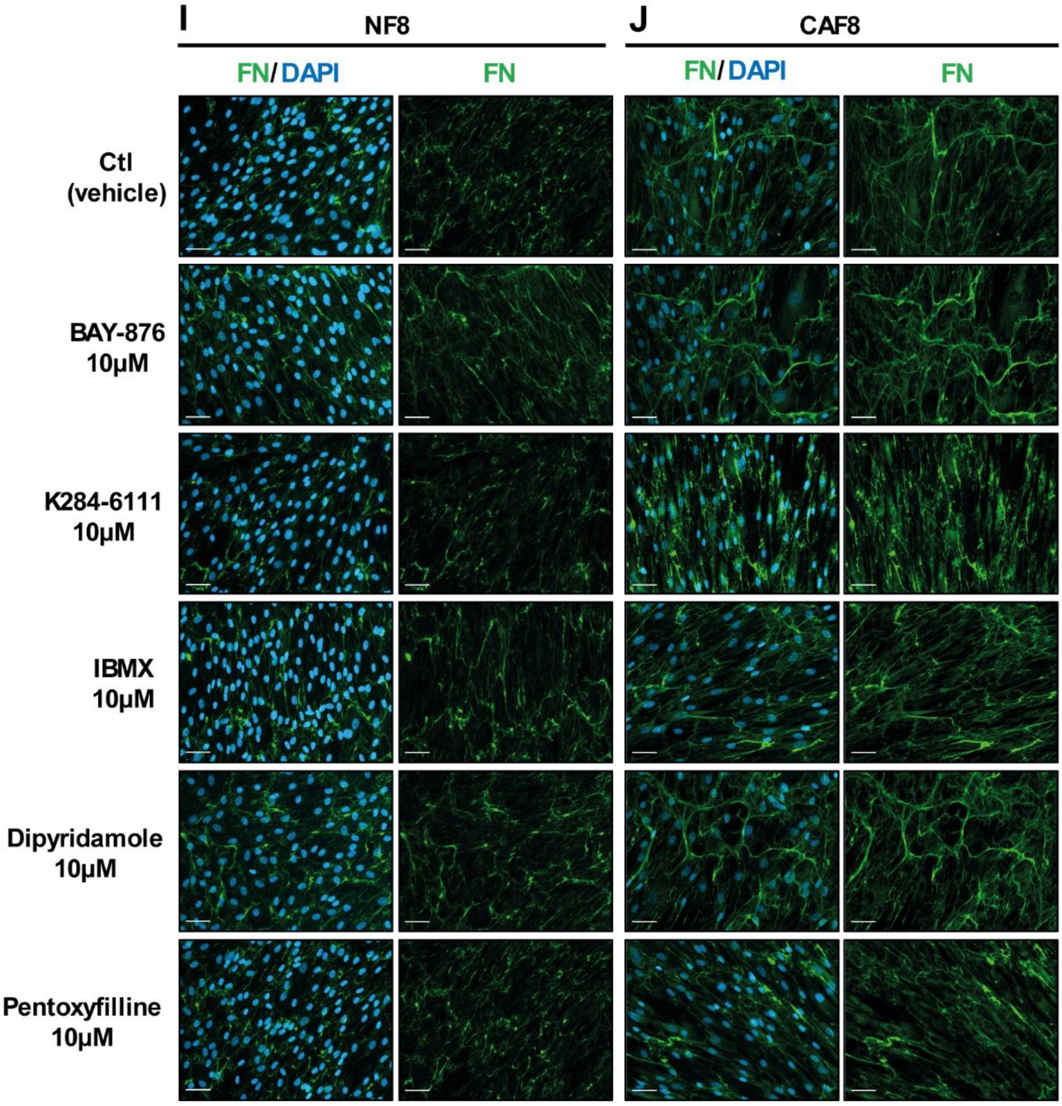


Figure continues next page ▶

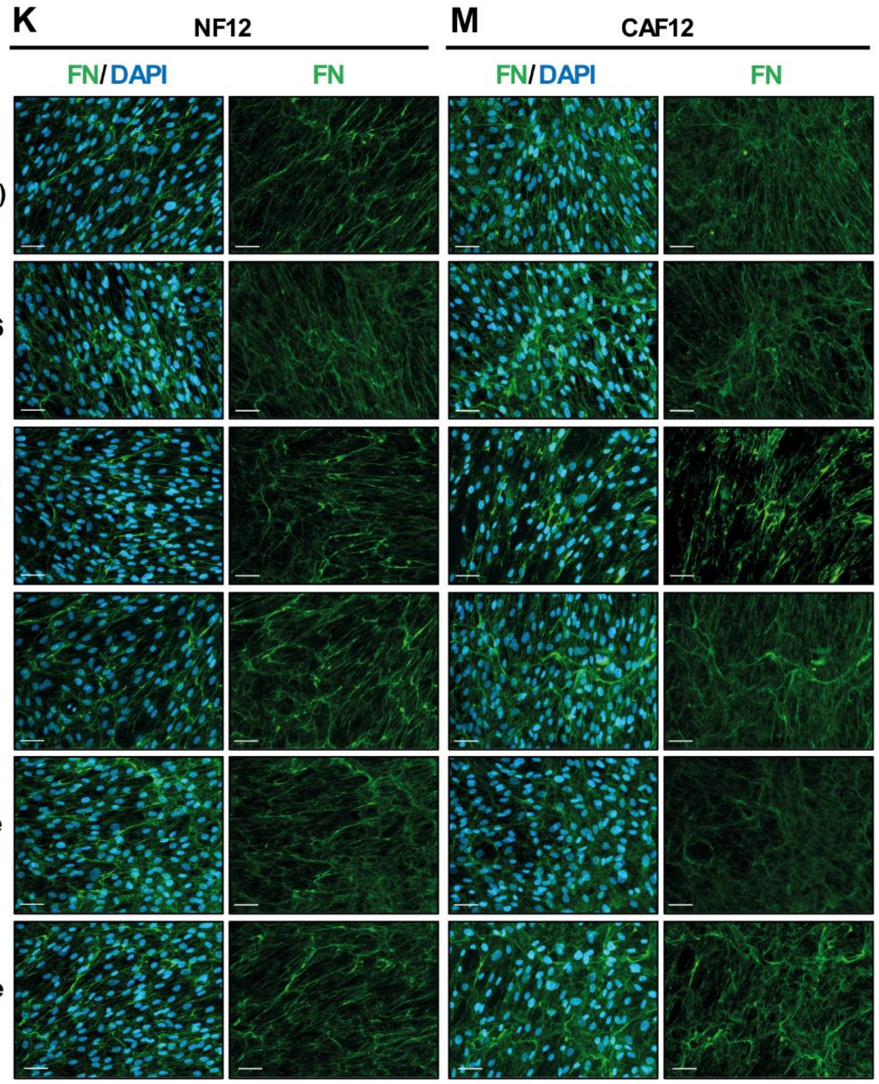


**Supplementary Figure S7. Effect of pharmacological inhibitors targeting CAF-related proteins on ECM synthesis and deposition.** Representative immunofluorescence images of fibronectin matrices derived from NF1 (**A**), CAF1 (**B**), NF2 (**C**), CAF2 (**D**), NF4 (**E**), CAF4 (**F**), NF7 (**G**), CAF7 (**H**), NF8 (**I**), CAF8 (**J**), NF12 (**K**), CAF12 (**M**) after treatment with either vehicle (Ctl) or the indicated inhibitors (10 µM). Green = fibronectin (FN); blue = DAPI. Scale bar = 50 µm.

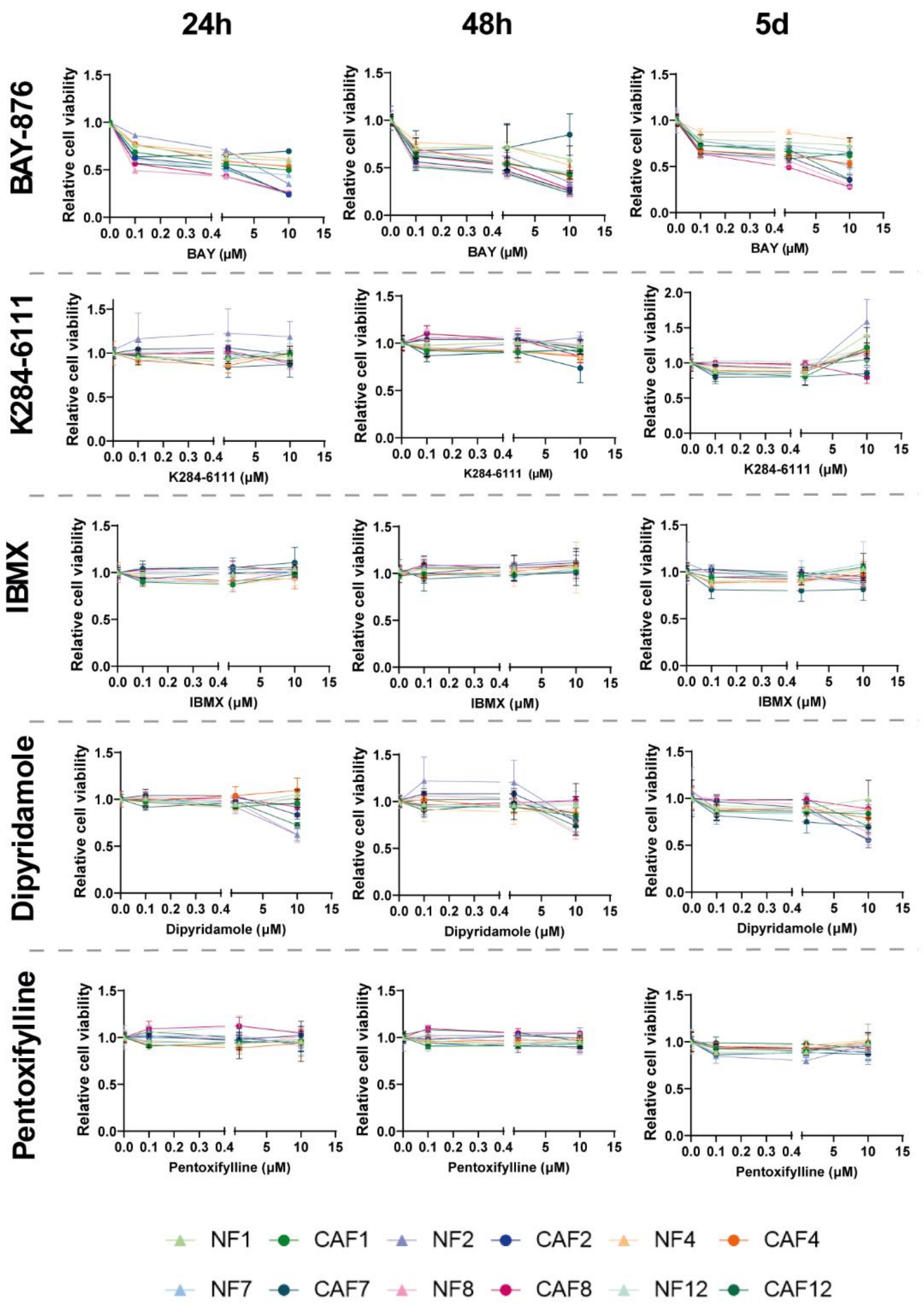


**Supplementary Figure S8**. **Effect of pharmacological inhibitors targeting CAF-related proteins on fibroblast cell viability**. Fibroblast viability normalized to day 0 and expressed relative to control condition (vehicle, DMSO) at 24 h, 48 h and 5 days following inhibitor treatment at 0.1, 1 and 10 μM. n = 2 independent experiments performed in triplicate.

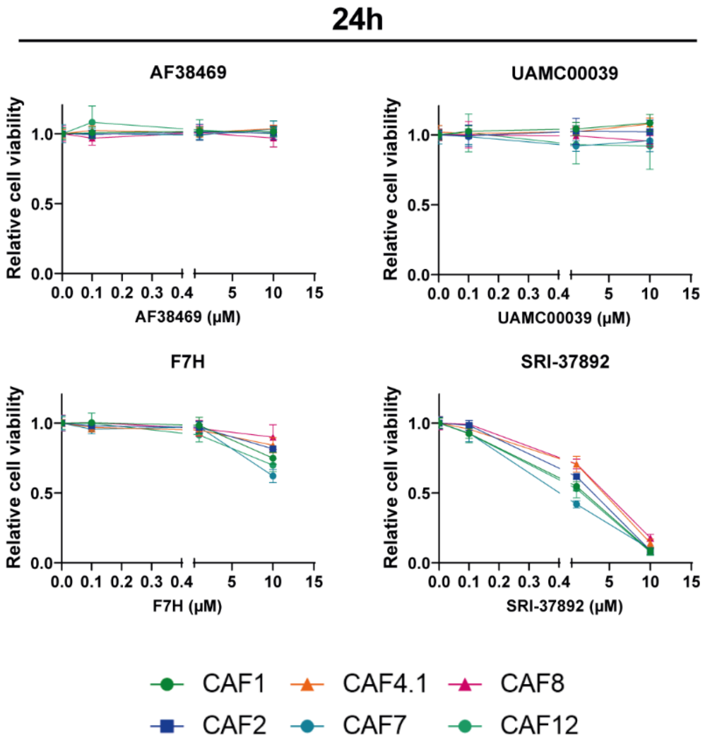


**Supplementary figure S9. Effect of pharmacological inhibitors targeting functional candidates on fibroblast viability.** The viability of the six primary CAF populations at 24 h was normalized to day 0 and expressed relative to control condition (vehicle) after treatment with the following inhibitors at 0.1, 1 and 10 µM: AF38469, UAMC00039, F7H and SRI-37892. n = 2 independent experiments in triplicate.
